# Co-infection of chickens with H9N2 and H7N9 avian influenza viruses leads to emergence of reassortant H9N9 virus with increased fitness for poultry and enhanced zoonotic potential

**DOI:** 10.1101/2021.04.05.438444

**Authors:** Sushant Bhat, Joe James, Jean-Remy Sadeyen, Sahar Mahmood, Holly J Everest, Pengxiang Chang, Sarah Walsh, Alexander MP Byrne, Benjamin Mollett, Fabian Lean, Joshua E. Sealy, Holly Shelton, Marek J Slomka, Sharon M Brookes, Munir Iqbal

## Abstract

An H7N9 low pathogenicity avian influenza virus (LPAIV) emerged through genetic reassortment between H9N2 and other LPAIVs circulating in birds in China. This virus causes inapparent clinical disease in chickens, but zoonotic transmission results in severe and fatal disease in humans. We evaluated the consequences of reassortment between the H7N9 and the contemporary H9N2 viruses of G1 lineage that are enzootic in poultry across the Indian sub-continent and the Middle East. Co-infection of chickens with these viruses resulted in emergence of novel reassortant H9N9 viruses carrying genes derived from both H9N2 and H7N9 viruses. These reassortant H9N9 viruses showed significantly increased replication fitness, enhanced pathogenicity in chicken embryos and the potential to transmit via contact among ferrets. Our study highlights that the co-circulation of H7N9 and H9N2 viruses could represent a threat for the generation of novel reassortant viruses with greater virulence in poultry and an increased zoonotic potential.

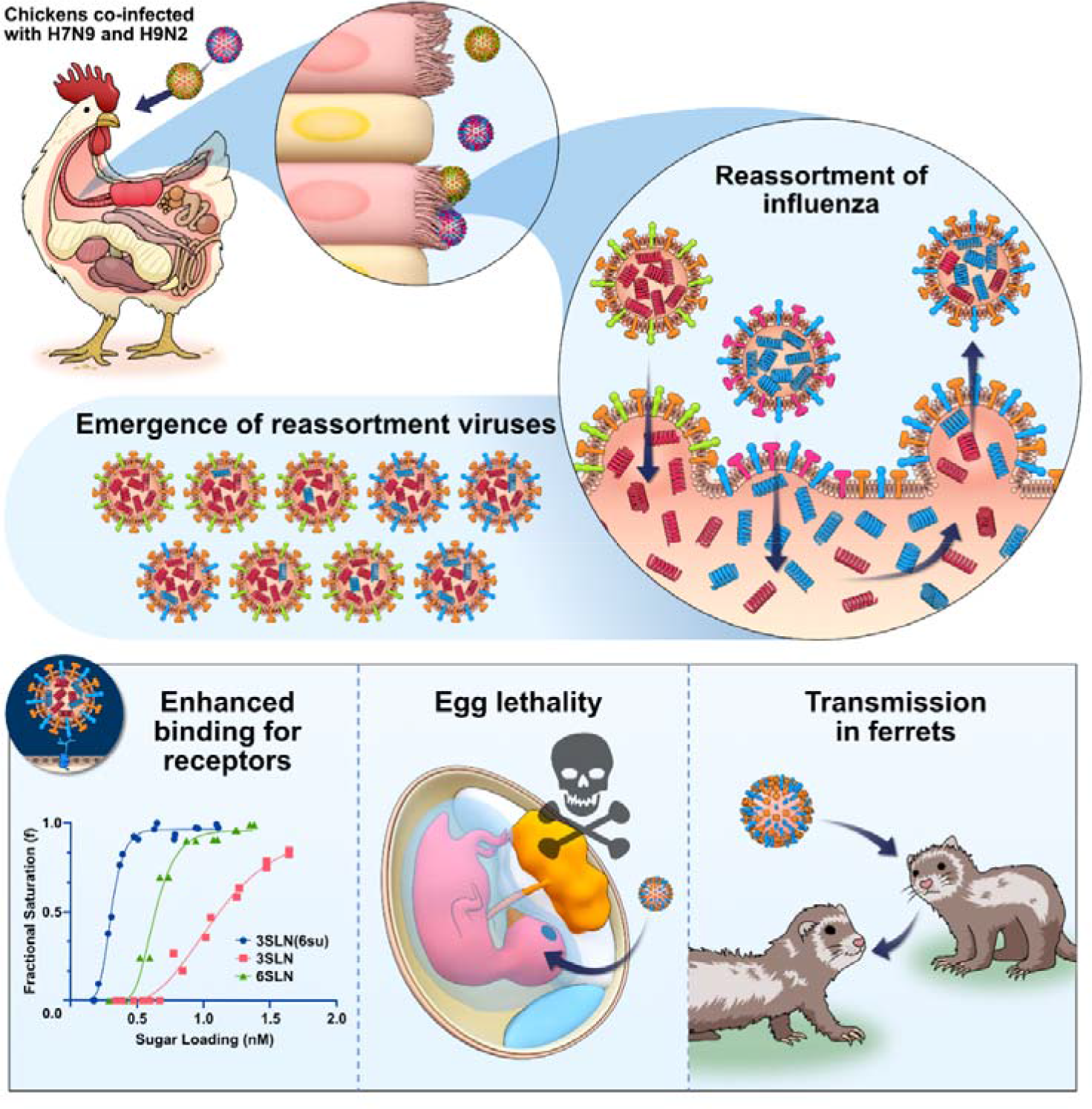
Graphical Abstract

**In Brief:** H9N2 viruses have a high propensity to reassort with other avian influenza viruses. We found that co-infection of chickens with H9N2 and H7N9 led to the emergence of reassortant viruses including the H9N9 subtype. Some reassortant H9N9 viruses exhibited increased replication fitness, increased pathogenicity in the chicken embryo, greater avidity for human and avian cell receptors, lower pH fusion and contact-transmission to ferrets. This study demonstrated the ability of viruses that already exist in nature to exchange genetic material, highlighting the potential emergence of viruses from these subtypes with increased zoonotic potential. There are nine H9 influenza A subtypes carrying different neuraminidase (NA) genes, including H9N9 viruses, while they are not common they do exist in nature as wildtypes (CDC).

**Highlights:**

- Co-infection of chickens with H7N9 and H9N2 led to emergence of reassortant H9N9 viruses
- Reassortant H9N9 viruses had an increased replication rate in avian and human cells
- Reassortant H9N9 viruses had a lower pH fusion and significantly higher receptor binding to α 2,3 sialoglycans
- Reassortant H9N9 replicated in ferrets at similar levels compared to H7N9 and transmitted via direct contact
- Ferrets exposed to reassortant H9N9 by aerosol contact were also found to be seropositive
- Experimental simulation of events that may occur naturally with circulating viruses has demonstrated the risk of emergence of viruses with increased zoonotic potential.

## INTRODUCTION

Novel human influenza A virus (IAV) infections during the past decade have included the H7N9 subtype, first isolated from humans in China, in early 2013 (Li et al., 2014). The virus was shown to be of avian origin, with zoonotic cases shown to be associated with exposure to infected birds at live poultry markets. Phylogenetic analyses revealed that the genotype of this avian influenza virus (AIV) arose naturally through complex genetic re-assortment events. The haemagglutinin (HA) gene was related to those identified in Eurasian-lineage H7N3 viruses found in ducks in Zhejiang, and its neuraminidase (NA) gene was closely related to that possessed by H7N9 viruses circulating in wild migratory birds in Korea (Gao et al., 2013). Furthermore, the six internal gene segments were most closely related to H9N2 viruses which are enzootic in chickens in China (Gao et al., 2013). This novel H7N9 virus was characterised as a low pathogenicity (LP) AIV (Kageyama et al., 2013) and caused mild or unapparent clinical disease in domestic chickens and ducks (Pantin-Jackwood et al., 2014), although a more severe pathogenesis may occur in turkeys (Slomka et al., 2018). Since 2013, transmission of this H7N9 virus from infected birds to humans has resulted in over 1500 infections in China with a case fatality rate of over 39% (FAO, 2021). Despite occasional reports of suspected nosocomial transmission (Fang et al., 2015), there has been no sustained spread of H7N9 between humans (Watanabe et al., 2014).

The continued enzootic circulation of H7N9 in poultry in China also resulted in the acquisition of polybasic amino acids at the cleavage site of the HA glycoprotein, a genetic hallmark of highly pathogenic (HP) AIVs (Chen et al., 2017; He, L. et al., 2018; Yang et al., 2017). The H7N9 HPAIV variants had an ability to cause up to 100% mortality in chickens (Tanikawa et al., 2019) and have resulted in at least 32 recorded human infections (FAO, 2021; Wang et al., 2017).

The evolutionary trend of H7N9 viruses and other zoonotic reassortant influenza viruses in nature inferred an increased reassortment propensity of H9N2 compared to other co-circulating AIV subtypes (Chen et al., 2014; Feng et al., 2013; Gao et al., 2013; Yang et al., 2015). This observation suggested that co-circulation of these viruses may predispose towards reassortment events to produce further genotypes with unknown disease risks to both poultry and humans. The continued circulation of H7N9 and H9N2 viruses with other AIV subtypes enzootic in farmed and wild bird populations has resulted in emergence of novel reassortant viruses with variable pathogenesis, along with the potential for mammalian adaptation and zoonotic transmission (Cui et al., 2014; Ge et al., 2018; He, J. et al., 2018; Lu et al., 2014; Shi et al., 2018; Wu et al., 2013; Wu et al., 2015).

Eurasian H9N2 AIVs have diversified into three main lineages (G1, BJ94 and Y438) which have themselves evolved to be characteristic of the geographical region that they occupy (Peacock et al., 2019; Slomka et al., 2013). Thus, H9N2 AIVs are enzootic in poultry in Asia, the Middle East and Africa (Alexander, 2007; Zhang et al., 2009) where they cause mild to severe morbidity and mortality in different avian species depending on the virus genotype (Chrzastek et al., 2018; Pu et al., 2017; Wang et al., 2018; Zhang et al., 2009; Zhu, R. et al., 2018). Like the emergence of novel genotype H7N9 LPAIVs, the G1 lineage H9N2 viruses currently circulating in the Indian subcontinent and the Middle East also include viruses which have undergone genetic reassortment with the internal gene segments from regional H7N3 HPAIVs (Ali et al., 2019; Chaudhry et al., 2015; Fallah Mehrabadi et al., 2019; Iqbal et al., 2009; Lee et al., 2016; Nagarajan et al., 2009; Shanmuganatham et al., 2014; Shanmuganatham et al., 2016). These reassortant H9N2 viruses have an increased zoonotic potential (Shanmuganatham et al., 2016) and have been reported to be more virulent and transmissible between poultry and terrestrial wild birds compared with their progenitors (Iqbal et al., 2013; Seiler et al., 2018).

To examine the credible natural reassortment scenario between H7N9 and G1 lineage H9N2 viruses, we performed experimental co-infection of chickens with A/Anhui/1/2013/H7N9 (Anhui/13) virus and A/Chicken/Pakistan/UDL-01/2008/H9N2 (UDL/08) virus. The genetic composition and phenotypic characteristics of the emergent reassortant viruses were analysed. An H9N9 reassortant was appropriately selected for infection using a ferret model to further ascertain any potential zoonotic characteristics.

## RESULTS

### H9 subtype viruses display enhanced viral shedding compared to the H7 subtype in co-infected chickens

To investigate the propensity for *in vivo* reassortment to occur between an H9N2 (UDL/08) and H7N9 (Anhui/13) virus in chickens, we needed to be able to identify the proportions of progeny viruses which contained the H7 or H9 HA gene segments. We developed an array of RT-qPCR assays, for each influenza gene segment, which could specifically detect, and discriminate the origin of the gene segment as either H9N2 (UDL/08) or H7N9 (Anhui/13) (Supplementary Table S1). Each gene segment specific assay yielded equivalent sensitivity and performance (Supplementary Figure S1) and were equivalent to the generic pan-avian influenza M-gene RT-qPCR (data not shown).

**Table 1:**
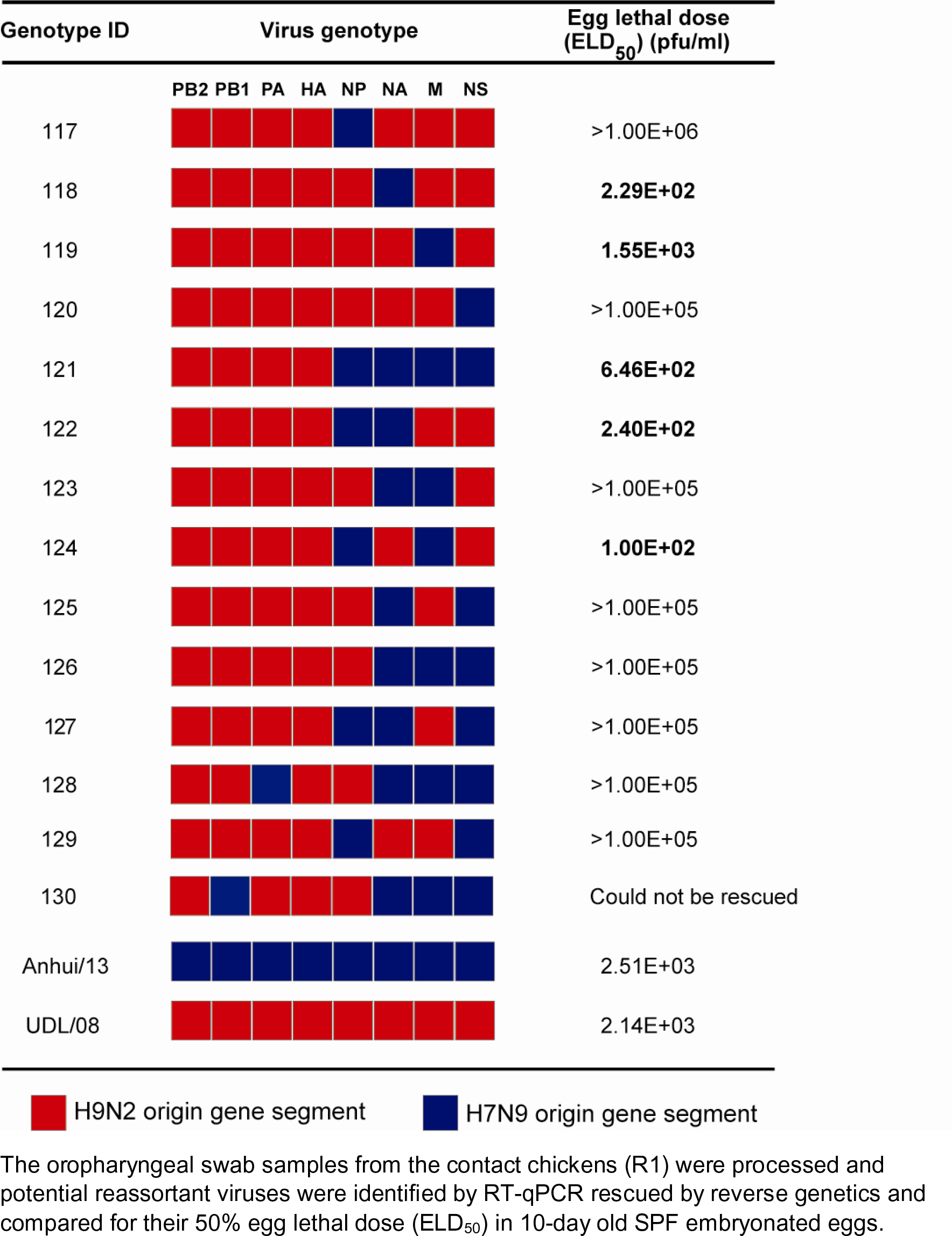
Genotypes of the potential reassortant viruses in swabs samples which emerged after co-infection and transmission to R1 chickens. The oropharyngeal swab samples from the contact chickens (R1) were processed and potential reassortant viruses were identified by RT-qPCR rescued by reverse genetics and compared for their 50% egg lethal dose (ELD_50_) in 10-day old SPF embryonated eggs.

Initially we used three AIV RT-qPCRs (H7-RT-qPCR, H9-RT-qPCR and the generic M-gene-RT-qPCR) to assess the oropharyngeal and cloacal shedding from chickens (i) directly co-infected with H7N9 and H9N2 (D0), (ii) chickens placed in direct contact with the co-infected chickens (R1), (iii) directly infected with H7N9 only, (iv) directly infected with H9N2 only (Figure 1). In the transmission experiment, the M-gene RT-qPCR detected positive viral RNA shedding, both oropharyngeal and cloacal swabs in all nine D0 (directly infected) and all nine R1 (contact) chickens (Figure 1A, C, E and G), thereby demonstrating successful infection of the D0 chickens and transmission to all nine R1 contacts. Oropharyngeal shedding of viral RNA had declined and resolved by 8-9 day post infection (dpi) for most of the chickens, except for # 41 where shedding was still detectable at 10 dpi (Figure 1A and C), and subsequently it was shown that all shedding had ceased by 11 dpi in all chickens (data not shown). Cloacal shedding of viral RNA however, was less prominent and at a generally lower titre in the nine D0 chickens (Figure 1E) compared to the oropharyngeal shedding, but higher, and of more sustained duration, for the R1 contacts (Figure 1G). All cloacal shedding in the D0 and R1 chickens had ceased by 9 dpi. Virus RNA shedding in the directly co-infected group (D0) kept for post-mortem (PM) analysis at 2 dpi and 4dpi showed viral RNA in the oropharyngeal samples while viral RNA could not be detected in the cloacal samples at 2dpi (Figure 1B and F).

**Figure 1.**
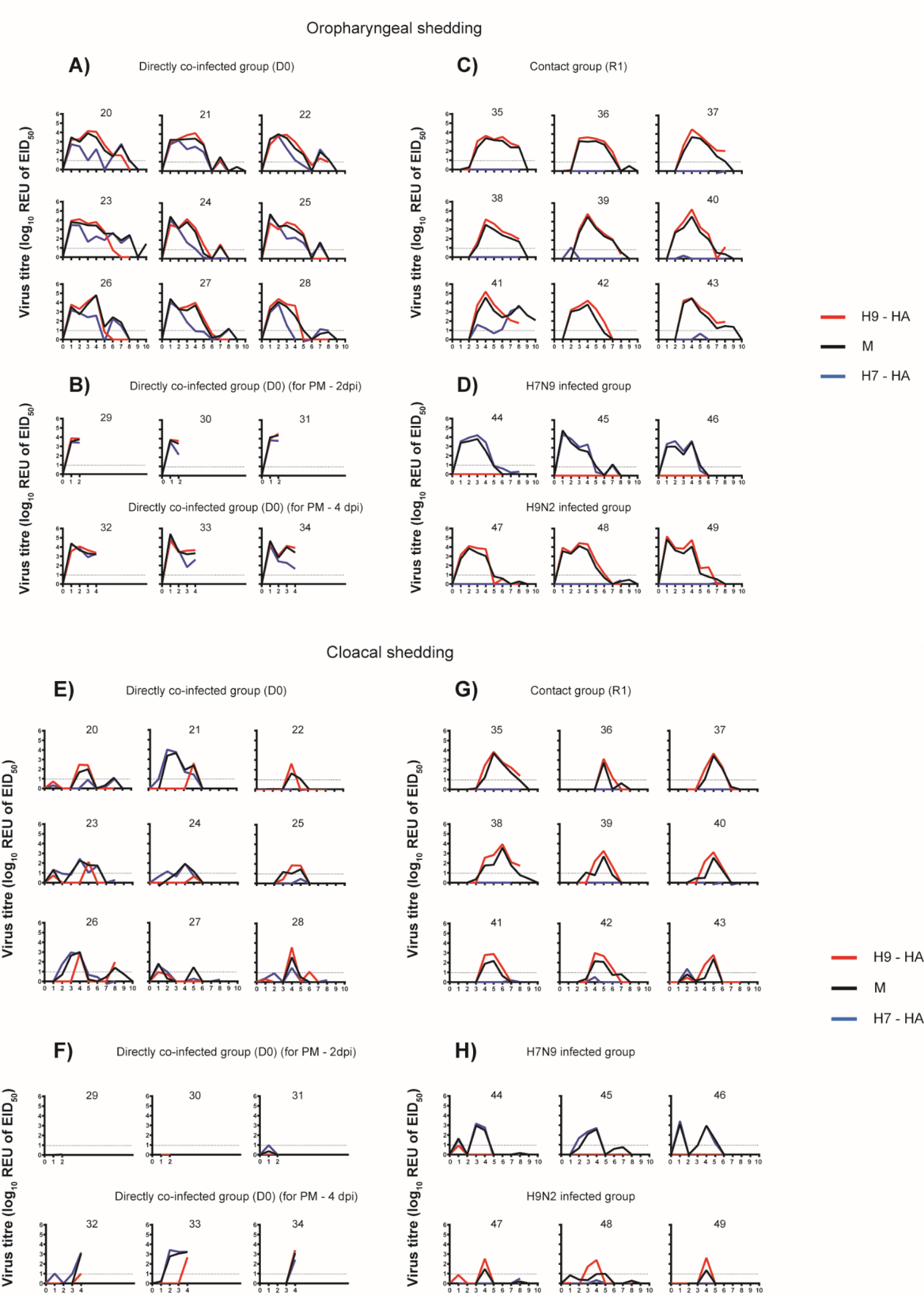
Detection of oropharyngeal (A-D) and cloacal (E-H) shedding, together with HA and NA genotyping to identify the respective viral subtypes which had replicated in chickens. Influenza A viral RNA shedding profiles expressed as relative equivalence units (REUs of EID_50_) (oropharyngeal and cloacal, as indicated) are shown for the individual infected chickens (numbered in individual panel). A standard curve was constructed using a dilution series of Anhui/13 viral RNA extracted form a known infectious titre (EID_50_/ml) of the virus, and tested by the M-gene, H7- and H9-specific RT-qPCRs (see Methods) which were shown to be equivalent in assay performance (Supplementary Figure S1). The Ct values were compared against an Anhui1/13 or UDL/08 RNA standards to determine relative equivalency units (REU of EID_50_). The dotted line represents the positive cut-off REU value. The chickens were directly co-infected with H9N2 UDL/08 virus and H7N9 Anhui/13 virus **(A and E)** to investigate transmission to contacts (**C** and **G**). Chickens were similarly co-infected and sacrificed at 2 and 4 dpi for virus dissemination in internal organs, with shedding similarly monitored (**B** and **F**). Shedding from chickens singly infected with either H7N9 Anhui/13 or H9N2 UDL/08 is also shown (**D** and **H**). The dotted horizontal line represents the REU of EID_50_ value at the limit of positive viral detection. Influenza virus shedding in all chickens had ceased by 11 dpi, so viral titres at subsequent swabbing days are not shown.

Measurement of viral RNA shedding by the H7- and H9-specific RT-qPCRs in the singly-infected chickens (Figure 1 D and H) showed the assays to yield very similar shedding results compared to the generic M-gene RT-qPCR, although some discrepancy was observed when shedding was at a low level. By applying the H7-and H9-specific RT-qPCR testing to the co-infected chickens, it was shown that, in the D0 chickens, the oropharyngeal shedding of RNA from both subtypes was initially of a similar magnitude, but at later days of shedding the level of H7 genome declined more rapidly while the shedding of the H9 subtype genome remained at higher levels, i.e., comparable to that detected by the M-gene RT-qPCR (Figure 1 A). The overall lower level of cloacal shedding from the D0 chickens suggested that H7 shedding either declined prior to the H9 shedding, or that H7 was weaker or below the positive threshold compared to the H9 shedding (Figure 1E). Among the R1 contact chickens, detection of viral RNA in the oropharyngeal and cloacal cavities was, however, exclusively of the H9 subtype (Figure 1C and G), the one exception being chicken # 41 where H7 oropharyngeal shedding appeared to increase after H9 shedding had declined, although cloacal shedding in the same chicken was due to the H9 subtype alone. In summary, while early shedding in the D0 chickens showed both subtypes to be detectable among the total AIV progeny, shedding of the H9 subtype endured for longer, with the H9 subtype being preferentially transmitted to infect the R1 chickens.

### Emergence of H9N2 and H9N9 as dominant subtypes in co-infected chickens

From the preliminary analysis with the H7, H9 and generic M-gene based RT-qPCRs, 4dpi was selected as the time point where oropharyngeal viral RNA appeared to be maximal or near maximal for the majority of the co-infected D0 and R1 chickens (Figure 1). Therefore, swabs from 4 dpi were selected to distinguish the origins (Anhui/13 or UDL/08) of all eight AIV genetic segments by using the segment-specific RT-qPCRs (Supplementary Table S1). Many of the 4 dpi swabs (oropharyngeal and cloacal) among the D0 chickens possessed a mixture of segments of both Anhui/13 and UDL/08 origins, although the UDL/08 origin H9 gene was strongly dominant among the HA segments analysed from the oropharyngeal swabs (Figure 2A). The Anhui/13 was restricted to only three of the D0 cloacal swabs (chicken #21, 23, 24), with no evidence of transmission to the R1 chickens (Figure 2B).

**Figure 2.**
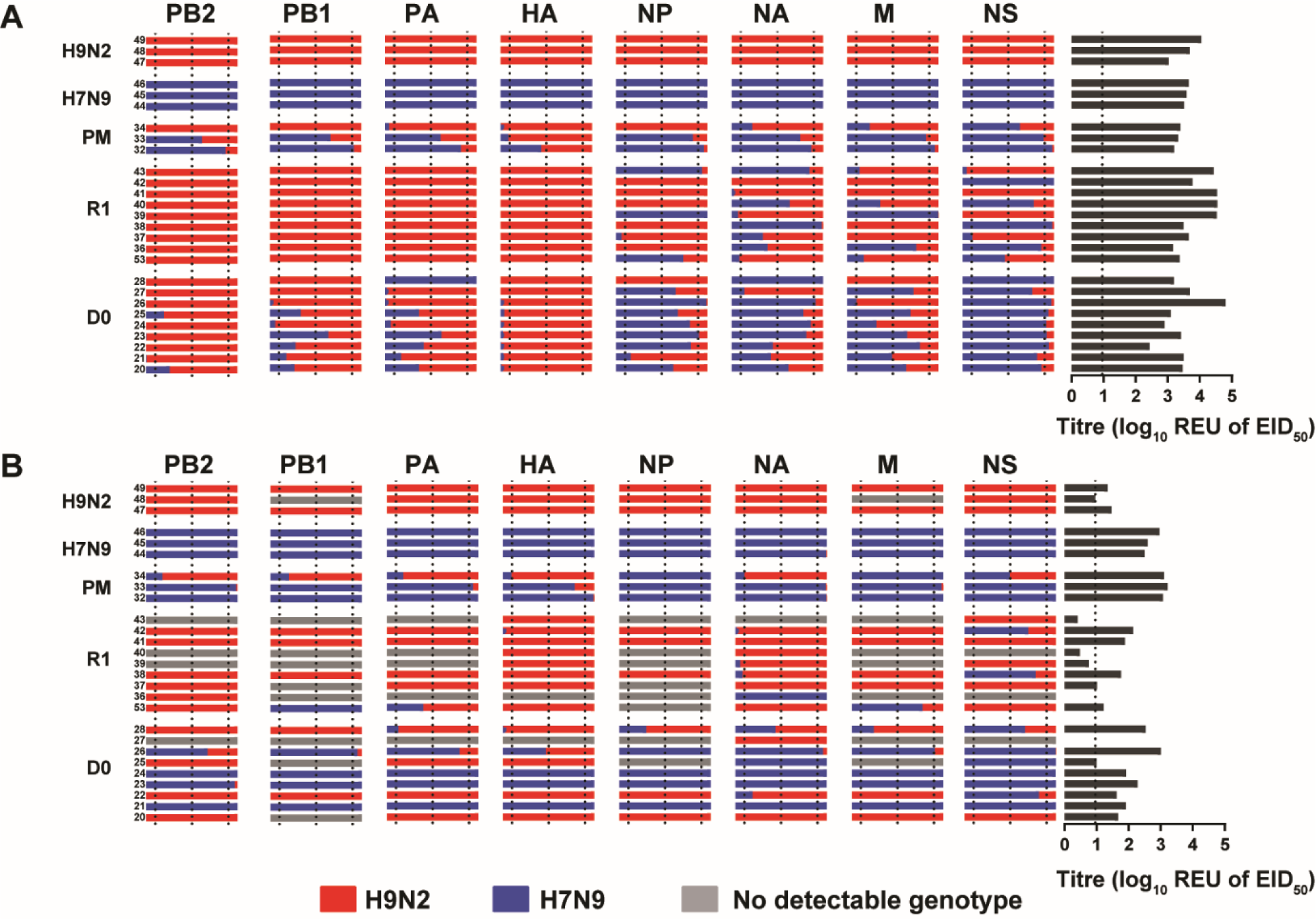
Genotyping of potential reassortant viruses shed from the oropharyngeal and cloacal cavities of D0 and R1 chickens at 4 dpi and 3 dpc, respectively. RNA extracted from oropharyngeal **(A)** and cloacal **(B)** swabs samples was used to quantify the proportions of each genetic segment using segment-specific RT-qPCRs for each parental virus strain (H9N2 UDL/08 shown in red; H7N9 Anhui/13 shown in blue). Swabs from individual chickens are represented in each horizontal row, with the proportions of each viral segment shown in separate columns. The Ct values were compared against an Anhui1/13 or UDL/08 RNA standards to determine relative equivalency units (REU of EID_50_). The REU values obtained from the segment-specific RT-qPCRs were converted to illustrate the percentage frequency of the origins of each gene, shown by the relative lengths of the horizontal red and blue bars. The vertical dotted lines within gene segment column represent 10%, 50% and 90% frequencies of each gene. Annotation on the left denotes the *in vivo* infected chickens: H9N2 and H7N9 correspond to the single-infected control groups; PM corresponds to the chickens which were pre-planned for cull and *post mortem* examination at 4 dpi (pathogenesis experiment); while D0 and R1 respectively indicate the direct- and contact-infected chickens following co-infection with both progenitor viruses; the final two digits of the individual chicken identifiers are discernible by the small font size at the left-end of each row. On the right, the AIV REU of EID_50_ for each chicken’s swab are shown by dark grey horizontal lines, with the broken vertical line indicating the REU positive cut-off. The failure to detect the origins of a given viral genetic segment (shown by grey horizontal bars) among several cloacal swabs tended to occur in those with low viral shedding values.

However, among the R1 chickens, contact transmission had resulted in an altered preponderance of UDL/08-origin segments, particularly for the HA (H9) and the three polymerase genes among the R1 oropharyngeal swabs where no corresponding Anhui/13- origin segments were detected at all (Figure 2A). These R1 oropharyngeal swabs included a variety of potential reassortants (genotypes), as evidenced by varying proportions of the mixed origins of the NP, NA, M and NS genes. Therefore, a mix of novel genotypes (which included the H9N2 and H9N9 subtypes) appear to have emerged (or were in the process of emerging) from the oropharynx of the R1 chickens. The reassortrant H7N2 subtype may have consisted of a small minority viral population at the D0 stage which failed to transmit to the R1 chickens (Figure 2). Overall, these results indicated a stronger replicative and transmissible fitness for the HA and polymerase gene segments of UDL/08 H9N2 virus origin.

### Virus dissemination in the D0 co-infected chickens

To address virus dissemination in chickens co-infected with H7N9 and H9N2 viruses, three chickens were sacrificed at 2 dpi and at 4 dpi. M-gene RT-qPCR revealed the highest viral RNA load in nasal turbinates (estimated as log_10_ REU of 50% Embryo Infectious Dose (EID_50_)) EID_50_) compared to all the tissues (Supplementary Figure S2). The viral RNA levels in the nasal turbinates were significantly higher at 2 dpi (1.85x10^4^ REU of EID_50_) compared to 4 dpi (1.91x10^3^ REU of EID_50_) (P<0.05), but there was no evidence of detectable infection in other organs within the respiratory tract (data not shown). The next highest viral loads were detected in the brain (1.08x10^3^ REU of EID_50_) followed by the cecal tonsils (3.43x10^2^ REU of EID_50_), but the difference was not significant between 2 dpi and 4 dpi (P>0.05), although RNA levels were higher in brain on 4 dpi. Several other organs revealed very low or sub-threshold levels (<1x10^1^ REU of EID_50_) of viral infection.

### Seroconversion in D0 co-infected chickens and the R1 contacts

Serum samples were collected at 14 dpi from co-infected (D0) and contact (R1) chickens and tested for antibodies by haemagglutination inhibition (HI) assay. The three single-infected control chickens seroconverted by 14 dpi to their respective homologous H7 or H9 subtype antigen (Figure 3). The nine co-infected D0 chickens showed a stronger seroconversion in the form of anti-H9 than anti-H7 antibodies at the same 14 dpi time-point. Eight of nine (89%) R1 contacts bled at the same time (13 day post-challenge (dpc)) reacted to only the H9 subtype antigen, with R1 chicken # 41 showed HI titre against H9 antigen but registered a weaker seroconversion against the H7 antigen (Figure 3). The viruses detected from this chicken (#41) included both the H7 and H9 HA segments during oropharyngeal shedding (Figure 1C).

**Figure 3.**
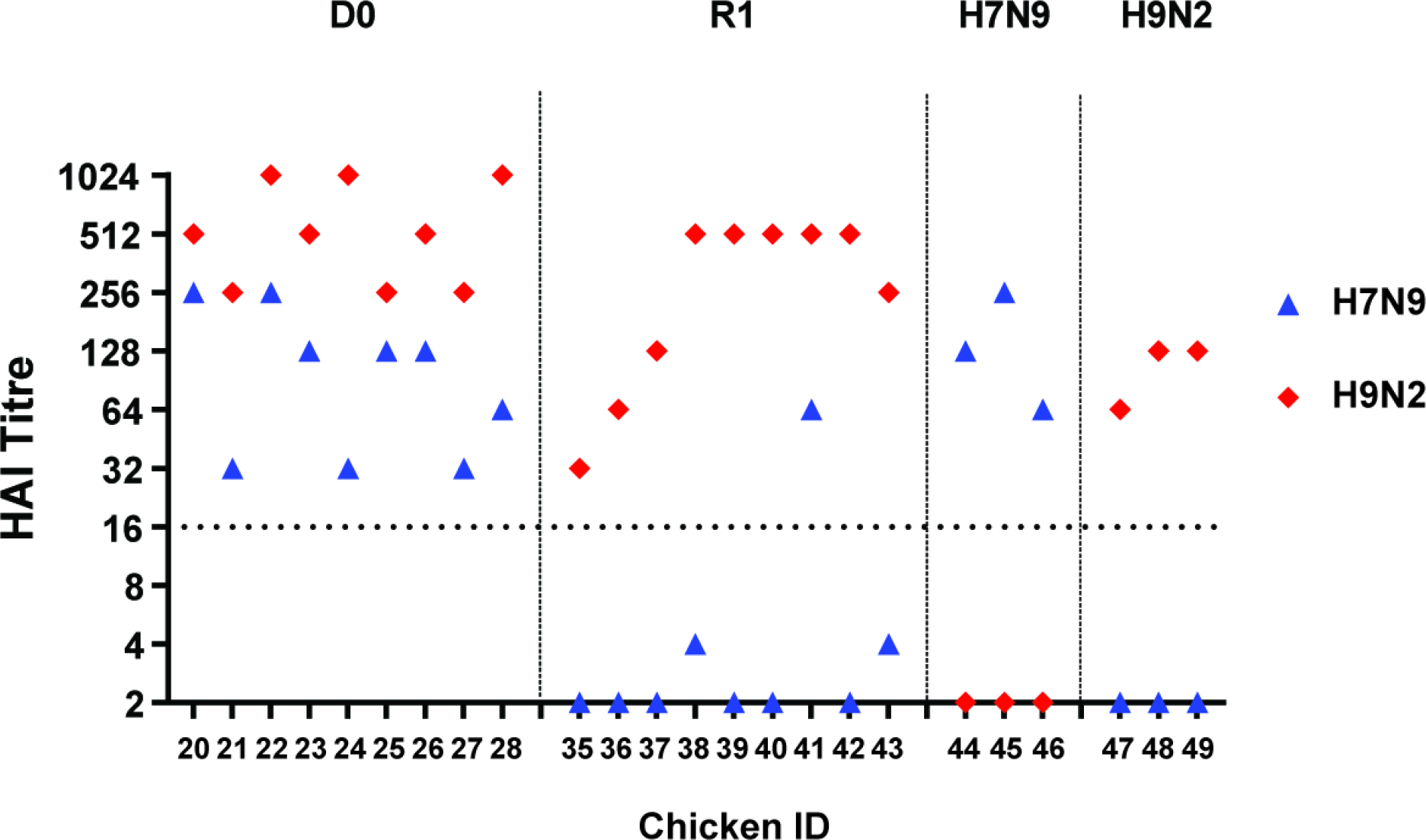
Seroconversion in chickens at 14 dpi / 13 dpc demonstrated by the haemagglutination inhibition (HI) test. Seroconversion in chickens directly co-infected (D0), co-infected contacts (R1) or singly-infected with H7N9 (Anhui/13) or H9N2 (UDL/08) viruses, as indicated by the headers. Symbols represent chicken HI titres against H7N9 (blue) and H9N2 (red) homologous antigens. The broken horizontal line indicates the HI positive cut-off value of 16 haemagglutination units. The 14 dpi time-point for D0 chickens corresponded to the 13 dpc time-point for the R1 chickens because the latter were introduced at 1 dpi.

### The ribonucleoprotein complex of H9N2 displays higher polymerase activity compared to that of H7N9 in chicken cells

An increase or decrease in polymerase activity can also be linked with the replication fitness or adaptability of a reassortant virus in a target host. As we previously observed enrichment of UDL/08 polymerase genes in the reassortant viruses, we therefore, investigated ribonucleoprotein (RNP) activity of polymerase gene segments of reassortant viruses using minireplicon assay. We quantified the polymerase activity of different RNP combinations obtained from the parental UDL/08 (H9N2) or Anhui/13 (H7N9) viruses. The RNP complex consisting of all four genes (PB2, PB1, PA and NP) from UDL/08 H9N2 produced more than 200% activity (P<0.0001) compared to Anhui/13 H7N9 RNP in chicken DF-1 cells at 39 ℃ (Figure 4A). By including Anhui/13-origin PB1 or PA on the UDL/08 background significantly reduced the activity compared to that of the unaltered UDL/08 RNP. However, inclusion of the Anhui/13-origin PB2 or NP produced greater polymerase activities compared to the unaltered UDL/08 (P<0.0001). These results showed that the RNP complex of UDL/08 has a greater polymerase activity compared to the RNP complex of Anhui/13 virus in DF-1 cells, with this greater activity attributed to the UDL/08 PB1 or PA gene segments.

**Figure 4.**
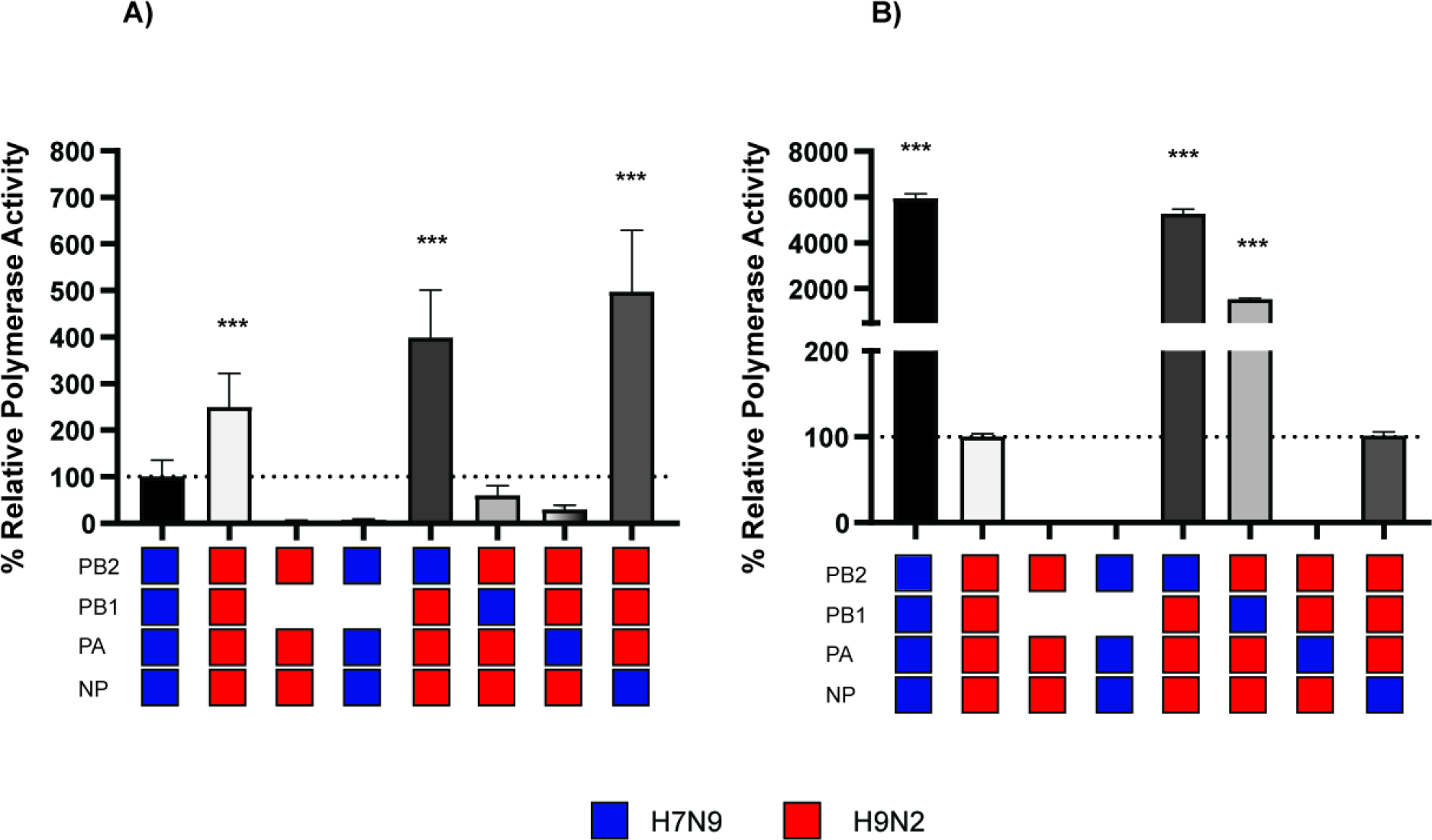
Minireplicon assay of the ribonucleoprotein (RNP) complexes of H7N9 Anhui/13 and H9N2 UDL/08 and four mixed RNP combinations. The RNP gene complexes derived from Anhui/13 and UDL/08 viruses were reconstituted by transfection of (A) chicken DF-1 and (B) human HEK-293T cells, along with four mixed RNP combinations from the H7N9 and / or H9N2 viruses. The cells were incubated at 37℃ (HEK-293T) or 39℃ (DF-1). At 24 hours post-transfection, cells were lysed and luciferase activities measured. Co-transfection of plasmids without PB1 served as a negative control for RNP activity. Percent polymerase activity was relative to that measured in the corresponding control H9N2 or H7N9 transfections. *** indicates P value <0.0001.

The RNP complex of UDL/08 H9N2 on the other hand showed a significantly lower polymerase activity compared to Anhui/13 H7N9 RNP in human HEK-293T cells (P<0.0001) (Figure 4B). Inclusion of PB2 and PB1 from Anhui/13 H7N9 on UDL/08 H9N2 background significantly increased the polymerase activity (P<0.0001) (Figure 4B). However, PA from Anhui/13 H7N9 reduced the polymerase activity and NP from Anhui/13 H7N9 resulted in a similar polymerase activity compared to UDL/08 H9N2. These results suggest that UDL/08 H9N2 virus polymerase genes are likely better adapted to avian hosts, whereas the Anhui/13 H7N9 virus polymerase genes are better adapted for humans.

### Multi-step replication kinetics and 50% egg lethal dose

Based on the RT-qPCR data (Figure 2), the gene segments contributed by parental UDL/08 H9N2 and Anhui/13 H7N9 towards the reassortant viruses which could have formed and shed from the oropharynx and cloaca of R1 chickens were identified and attempted for virus rescue by reverse genetics (RG). These included 11 genotypes (117-127) from the oropharyngeal swab samples and three (128, 129 and 130) from cloacal swabs (Table 1). The reassortant viruses and two parental strains were compared for their plaque forming ability (Supplementary Figure S3), chicken embryo lethality (Table 1) along with replication in primary chicken kidney cells (CK) (Figure 5), MDCK cells (Figure 6) and human A549 cells (Figure 7). For the replication of progenitor strains in CK cells, Anhui/13 H7N9 clearly displayed significantly greater kinetics than UDL/08 at 48 hrs post-infection (P<0.0001) (Figure 5n, but included in all panels). Acquisition of the M gene from Anhui/13 either alone (genotype 119) (Figure 5c), in combination with NP, NA and NS (genotype 121) (Figure 5e) or with NA and NS (genotype 126) (Figure 5j) gene segments from H7N9 substantially increased the replication of reassortant viruses at 48hrs post-infection compared to UDL/08.

**Figure 5.**
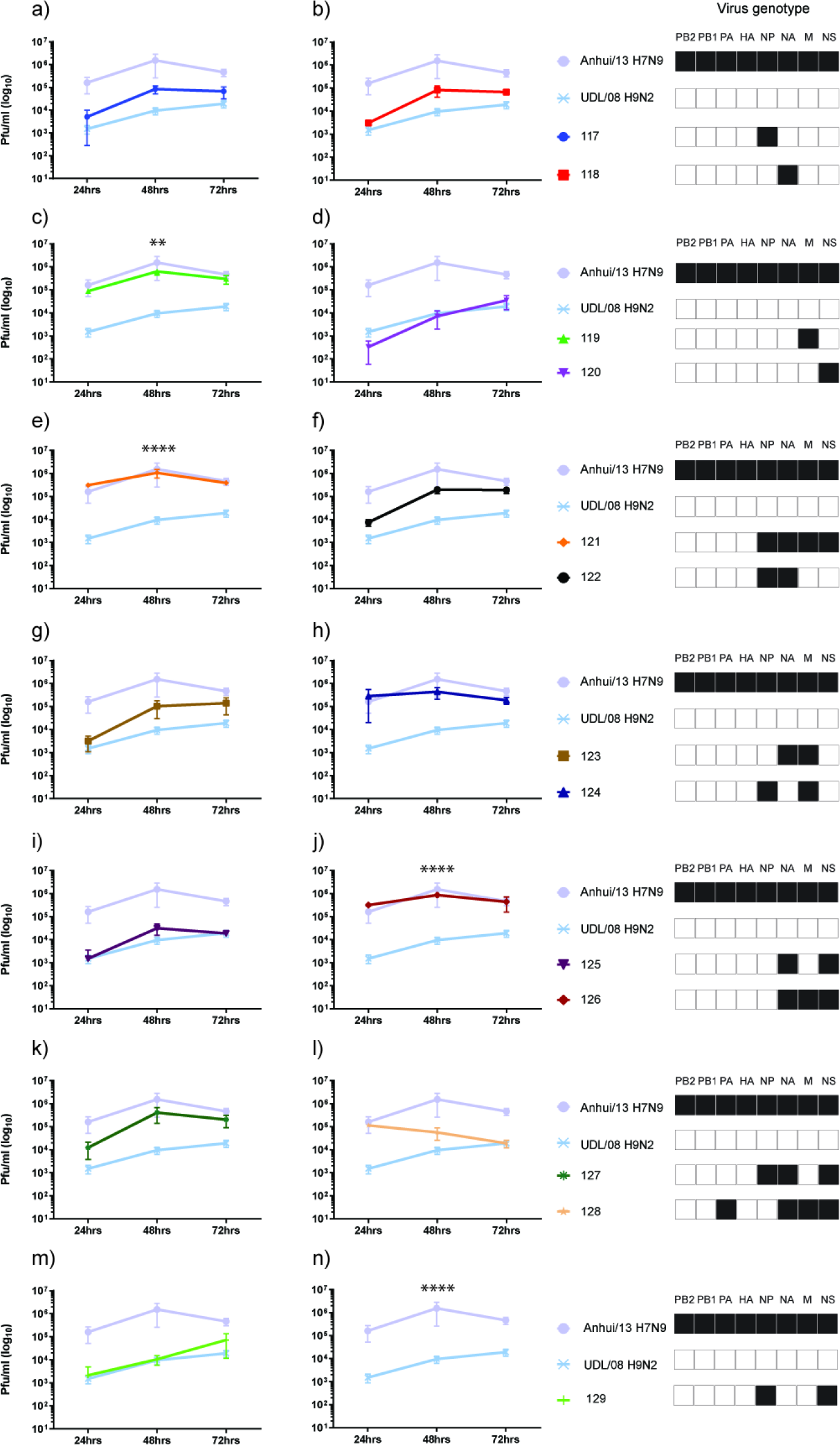
Multi-step replication kinetics of reassortant H9Nx viruses in primary chicken kidney (CK) cells. Primary CK cells were infected with 0.0002 multiplicity of infection (moi) of the 13 H9Nx reassortants or either of parental virus strains. Cell supernatants were harvested at 24hr, 48hr and 72 hr post-infection and titrated by plaque assay. Each time point corresponds to the mean of four biological replicates with standard deviations indicated. Replication kinetics of each reassortant virus compared to parental Anhui/13 H7N9 and UDL/08 H9N2 viruses is shown in panels (a) to (n). The genotype of each reassortant virus is shown as a combination of black and white coloured boxes with black indicating Anhui/13 H7N9 origin and white indicating UDL/08 H9N2 origin gene segments. ** denotes P value <0.005 and **** denotes P value <0.0001 compared to UDL08 H9N2.

**Figure 6.**
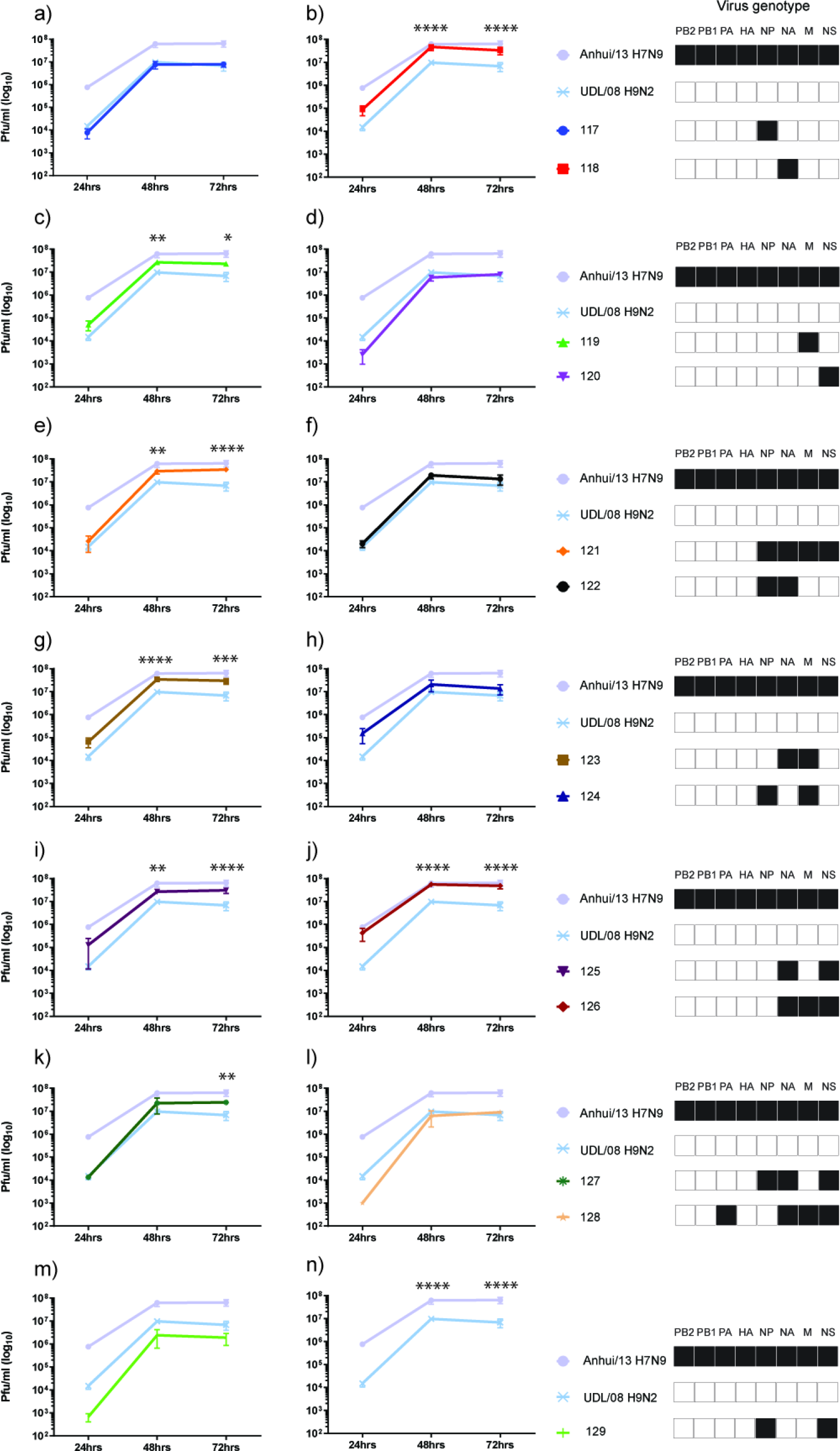
Multi-step replication kinetics of reassortant H9Nx viruses in Madin Darby Canine Kidney (MDCK) cells. MDCK cells (A) were infected with 0.0002 multiplicity of infection (moi) of the 13 H9Nx reassortants and both parental virus strains. Cell supernatants were harvested at 24hr, 48hr and 72 hr post-infection and titrated by plaque assay, and other annotations are as described in Fig 4. * denotes P value <0.05, ** denotes P value <0.005 and **** denotes P value <0.0001 compared to UDL/08 H9N2.

**Figure 7.**
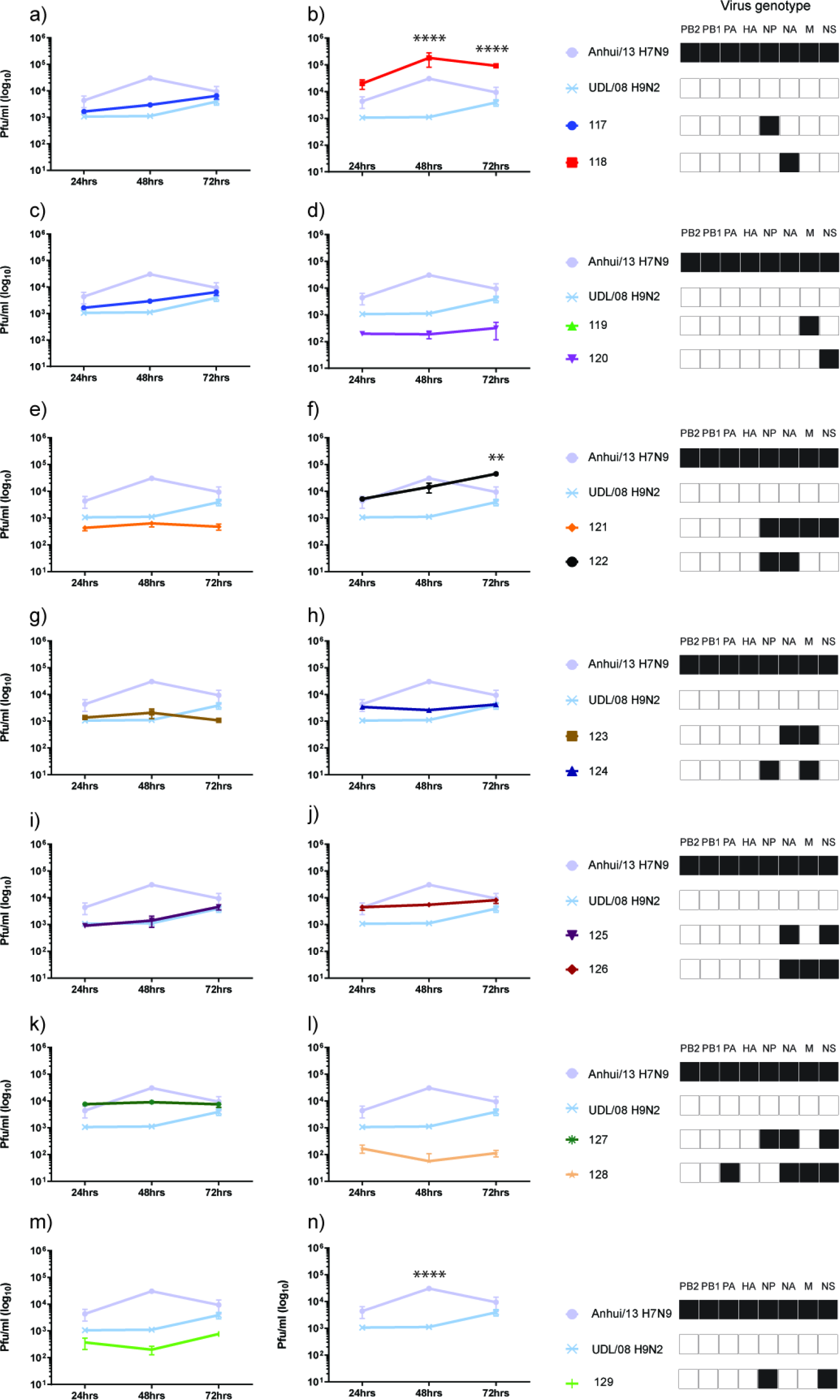
Multi-step replication kinetics of reassortant H9Nx viruses in human lung epithelial (A549) cells. A549 cells were infected with 0.05 multiplicity of infection (moi) of each reassortant and both parental virus strains. Cell supernatants were harvested at 24hr, 48hr and 72 hr post-infection and titrated by plaque assay, and other annotations are as described in Fig 4. ** denotes P value <0.005 and **** denotes P value <0.0001 compared to UDL/08 H9N2.

The viral replication kinetics in MDCK cells showed significantly higher replication for the Anhui/13 progenitor at 48 hrs and 72 hrs post-infection compared to the UDL/08 progenitor (P<0.0001) (Fig 6n, but included in all panels). Further, acquisition of the NA or M gene segment from H7N9 substantially increased the replication of reassortant genotypes 118 and 119, respectively, compared to UDL/08 H9N2 virus (Figure 6b, 6c). In human lung A549 cells, the replication kinetics of progenitor Anhui/13 was again significantly greater at 48 hrs post-infection compared to the UDL/08 progenitor (P<0.0001) (Figure 7n, but included in all panels). However, the reassortant virus with NA of Anhui/13 origin, namely genotype 118 (H9N9), showed significantly higher replication compared to both progenitor H7N9 and H9N2 viruses at 48 and 72 hrs post infection (P<0.0001) (Figure 7b). The reassortant H9N9 genotype 122 (having NP and NA of H7N9) also showed significantly higher replication compared to H7N9 and H9N2 viruses at 72 hrs post-infection (P<0.005) (Figure 7f).

Genotype 122 also displayed increased replication kinetics relative to UDL/08 in both CK and MDCK cells, albeit without statistical significance (P>0.05) (Figures 5f and 6f).

Five reassortant viruses also showed increased embryo lethality (lower embryo lethal dose (ELD_50_) compared to the progenitor viruses (Table 1). These five genotypes had acquired various segments from the Anhui/13 progenitor, namely (i) the genotype 118 H9N9 virus which acquired the NA gene; (ii) genotype 119 H9N2 virus which acquired the M gene; (iii) genotype121 H9N9 virus which acquired the NP, NA, M and NS; (iv) genotype 122 H9N9 virus which acquired the NP and NA and (v) genotype 124 H9N2 virus which acquired the NP and M. These observations suggest that the M and NA genes of the H7N9 virus enable greater adaptability for avian and mammalian hosts, respectively, as reflected in generally increased *in vitro* replication fitness.

### Plaque purification of viruses from R1 chicken swabs to identify viable reassortant viruses

Following initial co-infection of the D0 chickens, AIV RNA shedding at relatively high titres at 4 dpi (3 dpc) from the oropharyngeal cavity of the R1 chickens represented virus(es) that were sufficiently fit and had successfully transmitted within this host. To elucidate the exact constellation of genes within any viable reassortant virus(es), plaque purification of the oropharyngeal swab samples from all the nine R1 contact chickens was carried out in MDCK cells. Discrete plaques between a range of 16-28 were picked and RNA was extracted to fully characterise their genotype by segment specific RT-qPCRs. The genotype frequencies in each sample (expressed as percentage (%)) were calculated by taking the ratio of number of times a particular genotype appeared by total plaques isolated for a particular sample (Table 2). The genotypes identified by plaque purification reflected the overall genotyping as identified by RT-qPCR of swab samples (Figure 2). However, not all genotypes identified by RT-qPCR of swab samples could be isolated by plaque purification. The un-reassorted H9N2 UDL/08 was detected in four out of nine chickens between 100% to 13.6% genotype frequencies. In addition, a total of eight novel genotypes including single, double and triple segment reassortants were detected. Among all the viruses (Table 2) which had the highest genotype frequency (marked as *), included genotype 120 (7+1 reassortant H9N2 with NS from Anhui H7N9; isolated from four chickens at 100%, 12.5%, 4.2% and 3.7% genotype frequencies) followed by genotype 125 (6+2 reassortant H9N9 with NA + NS from Anhui H7N9; isolated from two chickens at 95.8% and 9.5% genotype frequency), genotype 122 (6+2 reassortant H9N9 with NP +NA from H7N9; isolated from two chickens at 88.9% and 12.5% genotype frequency) and genotype 124 (6+2 reassortant H9N2 with NP + M from H7N9; isolated from one chicken at 100% genotype frequency). UDL/08 H9N2 virus contributed substantially in terms of gene segments for different reassortant viruses and, interestingly, the H9N2 HA and polymerase gene segments (PB2, PB1, and PA) were conserved in 100% of the plaque isolates analysed in the R1 contact chickens (Table 2).

**Table 2:** Genotypes of the reassortant viruses isolated after plaque purification from the oropharyngeal swab samples from the co-infected contact chickens at 3 dpc. The genotype frequencies in each sample were calculated by taking the ratio of number of times a particular genotype appeared in each sample by total plaques isolated for a particular sample and expressed as percentage.

| Genotype ID | Chicken ID | % genotype frequency<br>(number of plaques) | Virus genotype |  |  |  |  |  |  |  |
| --- | --- | --- | --- | --- | --- | --- | --- | --- | --- | --- |
|  |  |  | PB2 | PB1 | PA | HA | NP | NA | M | NS |
| * UDL/08 | 41 | **100% (24)** |  |  |  |  |  |  |  |  |
|  | 40 | 66.7% (14) |  |  |  |  |  |  |  |  |
|  | 37 | 56.3% (9) |  |  |  |  |  |  |  |  |
|  | 36 | 13.6% (3) |  |  |  |  |  |  |  |  |
| * 120 | 42 | **100% (23)** |  |  |  |  |  |  |  |  |
|  | 37 | 12.5% (2) |  |  |  |  |  |  |  |  |
|  | 38 | 4.2% (1) |  |  |  |  |  |  |  |  |
|  | 43 | 3.7% (1) |  |  |  |  |  |  |  |  |
| 119 | 36 | 68.2% (15) |  |  |  |  |  |  |  |  |
|  | 43 | 7.4% (2) |  |  |  |  |  |  |  |  |
|  | 40 | 4.8% (1) |  |  |  |  |  |  |  |  |
| * 125 | 38 | **95.8% (23)** |  |  |  |  |  |  |  |  |
|  | 40 | 9.5% (2) |  |  |  |  |  |  |  |  |
| * 122 | 43 | **88.9% (24)** |  |  |  |  |  |  |  |  |
|  | 37 | 12.5% (2) |  |  |  |  |  |  |  |  |
| 118 | 40 | 19% (4) |  |  |  |  |  |  |  |  |
|  | 36 | 18.2% (4) |  |  |  |  |  |  |  |  |
| 117 | 37 | 12.5% (2) |  |  |  |  |  |  |  |  |
| * 124 | 39 | **100% (28)** |  |  |  |  |  |  |  |  |
| 127 | 37 | 6.3% (1) |  |  |  |  |  |  |  |  |

H9N2 origin gene segment H7N9 origin gene segment

Comparison of the percentage genotype frequency of reassortant viruses (Table 2) and their replication in human A549 cells (Figure 7), showed that genotype 122 notably had higher genotype frequency and showed increased replication. All the other reassortant genotypes which had a higher genotype frequency (genotypes 120, 124, 125) were attenuated for their replication in A549 cells. These results indicated that out of all the predominant genotypes, genotype 122 showed increased replication in human A549 cells. This observation warranted further investigation with respect to the zoonotic potential of genotype 122 H9N9 virus.

### Reassortant H9N9 virus has preferential receptor binding with α2,6 sialic acid receptor analogues compared to the parental UDL/08 (H9N2 virus)

Host receptor binding preference of influenza viruses is a critical determinant of host adaptation and airborne transmission in ferrets (Tumpey et al., 2007). The receptor binding specificity of parental and reassortant H9N9 viruses were quantified with synthetic sialoglycopolymers - α2,6 sialyllactosamine (6SLN), α2,3 sialyllactosamine (3SLN) or 1-4(6-HSO) GlcNAc (3SLN(6-su)), receptor analogues using bio-layerNeu5Ac α2,3Gal β1-4(6-HSO­_3_) GlcNAc (3SLN(6-su)), receptor analogues using bio-layer interferometry (Xiong et al., 2013). The selected reassortant H9N9 virus (genotype 122, Figure 8A) showed strong binding for 6SLN receptors which was comparable to that of Anhui/13 (Figure 8C). By contrast, the parental UDL/08 displayed only marginal and undetectable binding to 6SLN receptor analogues (Figure 8B). The reassortant H9N9 viruses also bound strongly to 3SLN compared to the parental UDL/08 H9N2 virus, which showed no binding (Figure 8A and B). The binding avidity for 3SLN (6-Su) was also stronger for the reassortant H9N9 viruses compared to UDL/08 H9N2 virus. These observations showed that genotype 122 H9N9 virus has increased receptor binding preferences for both avian and mammalian hosts compared to the genetic donor viruses.

**Figure 8.**
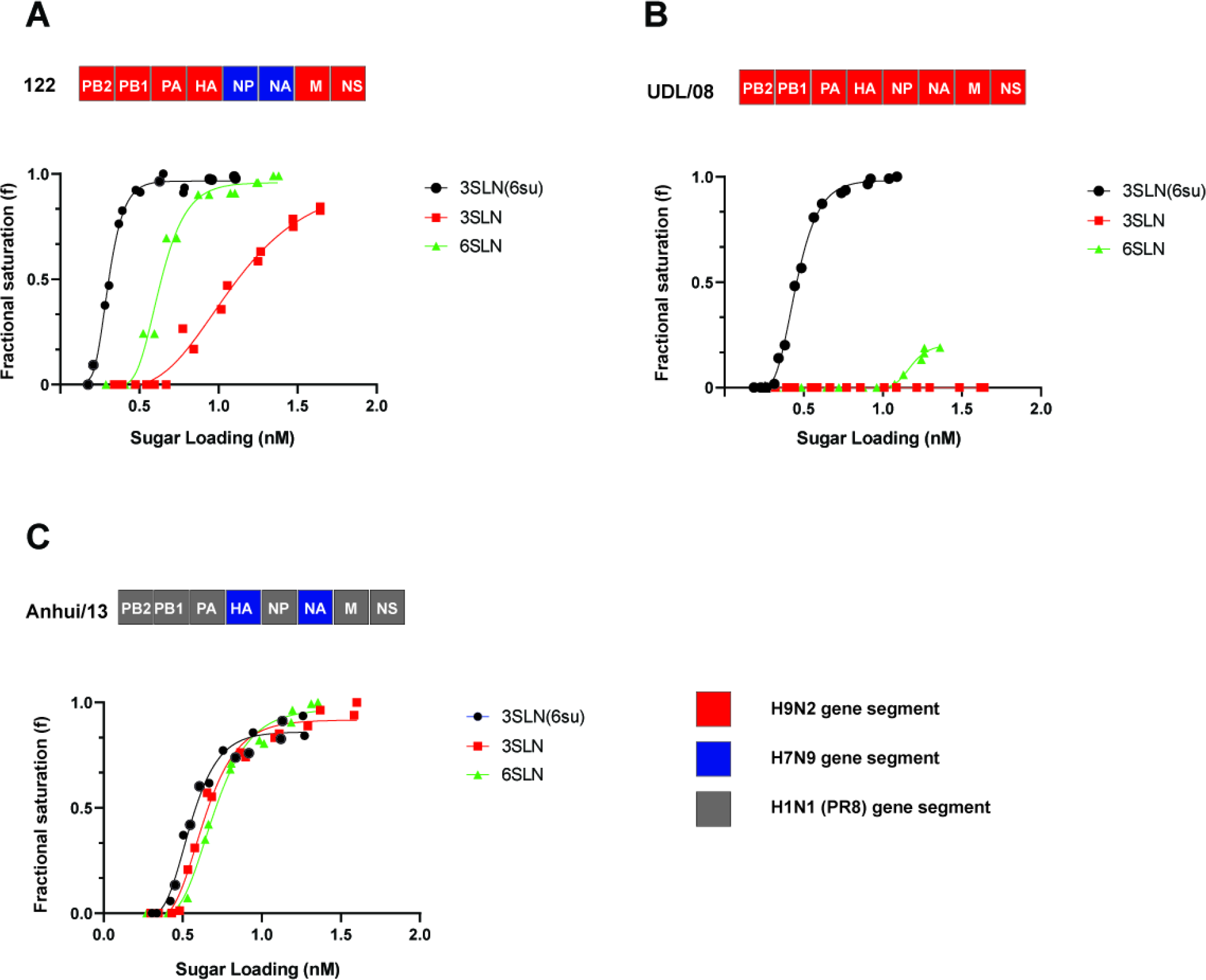
Receptor binding profiles of reassortant H9N9 virus compared to H9N2 and H7N9 viruses. A) Binding of 6+2 reassortant H9N9; genotype 122 [two genes (NP and NA) from Anhui/13 H7N9 and 6 genes from UDL/08 H9N2] to α-2,3-linked (3′SLN 6-sulphated) (black), α- 2,3-linked (3′SLN) (red), or α-2,6-linked (6′SLN) (green) sialylglycan receptors was determined by biolayer interferometry. Similar receptor binding profiles were determined for **(B)** UDL/08 H9N2 and **(C)** 2+6 reassortant H7N9 [2 genes (HA and NA) from Anhui/13 H7N9) plus 6 genes from PR8]. *Since interferometry involved testing of infectious virus, due to biosafety reasons the receptor binding of H7N9 was carried out using the 2+6 reassortant of H7N9 which included internal genes from PR8. * Since biolayer interferometry involved testing of infectious virus, due to biosafety reasons the receptor binding of H7N9 was carried out using the 2+6 reassortant of H7N9 which included internal genes from PR8

### Fusion pH of reassortant H9N9 virus compared to parental Anhui/13

The pH of fusion critically influences stability and infectivity of virus in the target host species. The viruses that are stable at lower pH carry greater propensity to retain infectivity in the human airway epithelium. We determined the fusion pH of the reassortant H9N9 virus (genotype 122) and the parental H7N9 and H9N2 viruses in Vero cells using a syncytium-formation assays. For H7N9 virus infections, cells showed optimal pH fusion at 5.6, while for reassortant H9N9 virus (genotype 122) and the parental H9N2 virus infected cells demonstrated optimal fusion at pH 5.4. This observation showed that the reassortant H9N9 virus has a greater pH stability of the HA compared to that of the H7N9 viruses.

### Assessment of the zoonotic risk of the selected reassortant H9N9 virus

We further assessed the zoonotic potential of the novel H9N9 genotype using ferrets as an animal model of infection in humans. The findings from the quantitative investigations of the viruses shed from the oropharynx of chickens along with the replication kinetics, receptor binding and fusion pH guided the assessment of genotype 122 (H9N9 reassortant) and its comparison to Anhui/13 H7N9 virus in a ferret transmission study. The ferrets directly-infected (D0) with Anhui/13 H7N9 were positive for virus RNA detected in the nasal wash samples from 2-10 dpi, while the D0 ferrets infected with H9N9 showed positive shedding from 2-6 dpi (Figure 9A and B). However, the peak level virus shedding was comparable in both D0 groups. Both the Anhui/13 H7N9 virus and reassortant H9N9 (genotype 122) virus exhibited 100% transmission efficiency from ferret-to-ferret when in direct contact (R1_DC_); all R1_DC_ contact ferrets became infected and shed virus from the nasal cavity (Figure 9A and B). However, ferrets sharing the same airspace but separated physically (indirect) via a dividing mesh in adjacent cages did not show detectable viral shedding in either groups (R1_In_).

**Figure 9.**
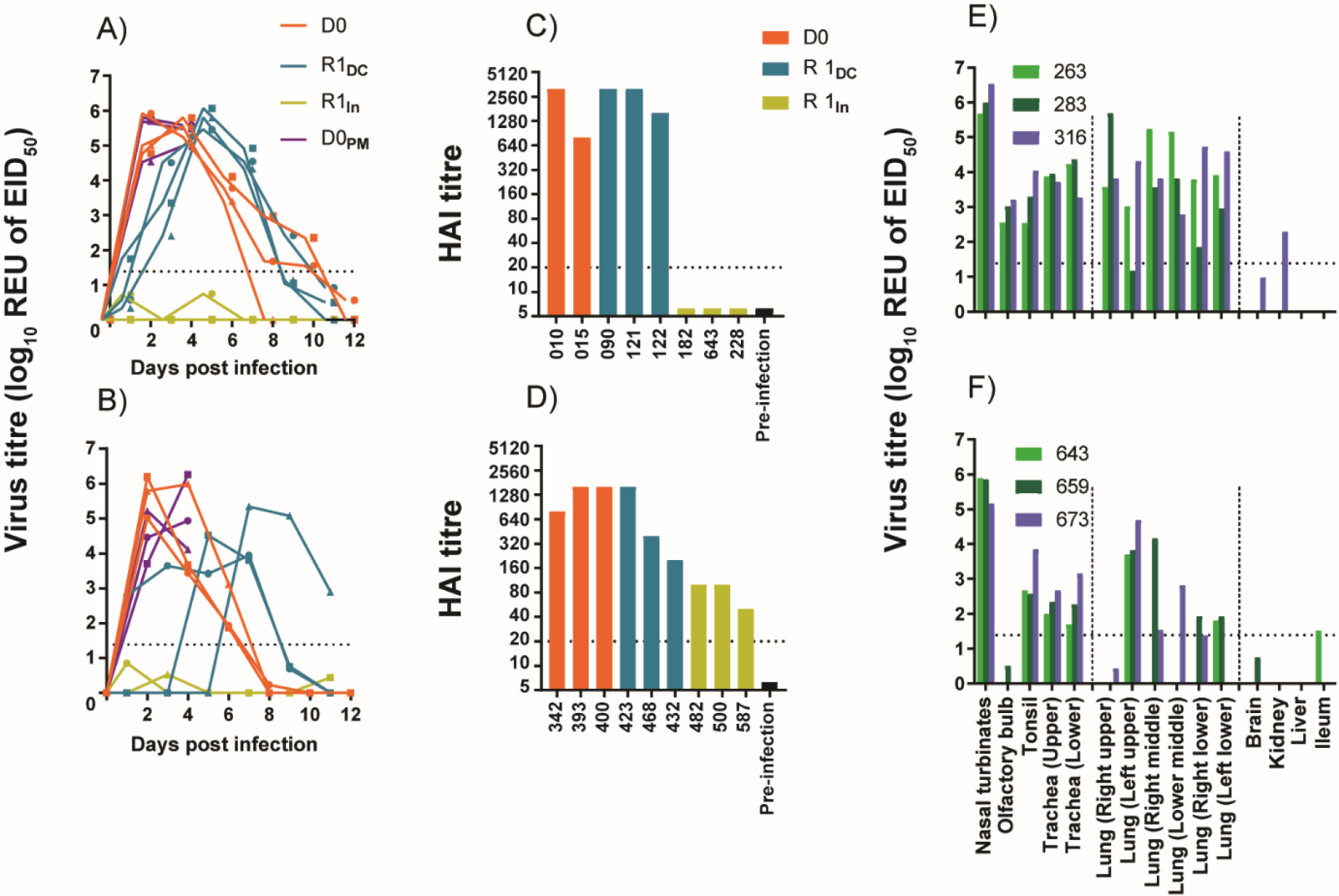
Detection of influenza A virus RNA and seroconversion in ferrets after intranasal inoculation with H7N9 and reassortant H9N9 viruses. Two separately housed groups of ferrets (D0, n=6 per group) were infected directly via the intranasal route with **(A)** Anhui/13 and **(B)** the reassortant H9N9 (genotype # 122). At 1 dpi, a group of ferrets (n=3) were placed in direct contact (R1_DC_) with each group of D0 ferrets, while another indirect contact group (R1_In_) of ferrets (n=3) were placed in an adjacent cage. Nasal washes were collected on alternate days from all ferrets which remained in the study until 12 dpi to determine viral shedding (REUs of EID_50_) by the M-gene RT-qPCR. Along the horizontal axis, dpi and dpc corresponds to time-points for the D0 and all the contact ferrets respectively. **(C, D)** Seroconversion in D0, R1_DC_ and R1_In_ ferrets at 14 dpi / 13 dpc. **(E, F).** Three ferrets from each D0_PM_ group were euthanized for postmortem on 4 dpi to assess virus dissemination in internal organs by M-gene RT-qPCR. The broken horizontal line corresponds to the positive cut-off value for the M-gene RT-qPCR (A, B, E and F) and the HI (C and D) tests.

All directly infected (D0) and direct contact (R1_DC_) ferrets seroconverted when tested by HI assay against homologous viruses (Figure 9C and D). None of the ferrets indirectly exposed to the Anhui/13 H7N9 virus infected group (R1_In_) seroconverted to H7N9 virus, but all ferrets indirectly exposed to the reassortant H9N9 virus infected ferrets (R1_In_) seroconverted to H9N9 virus (Figure 9C and D).

Three ferrets in each of the D0 groups were culled at 4 dpi in order to provide a range of organs, mainly from the respiratory tract, for post mortem examination. Both the Anhui/13 and reassortant H9N9 viral RNA were detected at high levels (>5log_10_ REU of EID_50_) in the nasal turbinates of the D0 ferrets, with the former also detected in the olfactory lobe (Figure 9E and F). Viral nucleoprotein was detected in the respiratory and olfactory epithelium of the nasal turbinates in all three infected ferrets (Supplementary Figure S4). In addition, histological lesions were identified in the respiratory epithelium of all ferrets for both viruses, and to a lesser extent in the olfactory epithelium for both viruses (Supplementary Figure S4). Viral RNA for both viruses was detected in the upper and lower trachea, however viral nucleoprotein antigen was not detected by IHC in the trachea in the H9N9 infected group (Supplementary Table S2). Anhui/13 virus RNA was detected in the upper, middle and lower lung tissues whereas the reassortant H9N9 virus RNA was higher in the upper left lobe of the lungs, with sporadic detection in the other lobes (Figure 9E and F and Supplementary Table S2). Overall, H7N9 infected ferrets showed more pulmonary lesions compared to H9N9 infected ferrets (Supplementary Figure S5). Neither of the H7N9 or H9N9 viruses was detected in the brain or liver, although Anhui/13 was detected at a low level (>2log10 REU of EID_50_) in the kidney of one infected ferret (Figure 9E and F), while H9N9 was detected in the ileum of one ferret at the limit of detection.

As regards to clinical changes, the D0 ferrets directly-infected with H9N9 reassortant experienced a negligible increase in body temperature and a modest reduction in weight (around 2-3%) at 2 dpi (Supplementary Figure S6 C and D), compared to Anhui/13 D0 infected ferrets which developed fever from 1 to 3 dpi and exhibited weight loss from 1 till 7dpi, which decreased to ∼10% of their starting weight in some ferrets (Supplementary Figure S6 A and B). The increase in body temperature correlated with peak viral shedding of Anhui/13 from the D0 ferrets (Figure 9A).

The ferrets placed in contact with the directly infected Anhui/13 and reassortant H9N9 infected ferrets did not develop a significant increase in body temperature or a weight loss.

## DISCUSSION

The novel H7N9 LPAIV (Anhui/13) emerged in 2013 in China through a triple reassortment event, producing a virus with all of the six internal genes derived from the G57 lineage (genotype S) of H9N2 viruses (Pu et al., 2015). Although the H7N9 virus came to prominence through its zoonotic phenotype, it is essentially an avian influenza virus (AIV) which has been circulating in avian species in China for several years (Ke et al., 2014). Co-circulation of H7N9 with several other AIV subtypes enzootic in birds in China, including H9N2, has resulted in extensive genetic reassortment that has led to the emergence of diverse H7N9 genotypes (OIE, 2018b; Su et al., 2018) which continue to infect humans and birds (Lam et al., 2015; Lu et al., 2014; Zhu, W. et al., 2018). H9 viruses have also reassorted with diverse NA subtypes leading to nine (N1-N9) known subtypes including H9N9 viruses (CDC).

As both H7N9 and H9N2 virus circulate naturally in avian population it is of value to determine what novel reassortants may emerge because of natural co-infection with these two virus subtypes. We investigated the reassortment potential of H7N9 virus with the G1- lineage H9N2 viruses that are enzootic in poultry in China and neighbouring countries (Biswas et al., 2018; Iqbal et al., 2009; Shanmuganatham et al., 2014). The experimental co-infection of chickens with H7N9 Anhui/13 and a G1-lineage H9N2 UDL/08 virus resulted in the emergence of reassortant influenza A viruses (IAVs) that efficiently transmitted to contact naive chickens. Many studies have investigated the generation of reassortant influenza A viruses in different hosts like chickens (Li et al., 2018), mallards, guinea pigs (Tao et al., 2014) (Ganti et al., 2021), swine (Zhang et al., 2018) and embryonated chicken eggs (Naguib et al., 2017), to the best of our knowledge, this is the first study illustrating co-infection of chickens with H7N9 and H9N2 IAVs resulting in emergence of reassortant H9N9 influenza viruses with a zoonotic potential.

For two influenza viruses to reassort most efficiently in a host, the viruses must establish a state of co-infection (Lowen, 2018), for which, productive infection of hosts is required (Fonville et al., 2015). In view of previous studies of experimental chicken infections where a high dose was indicated for H7N9 Anhui/13 (Slomka et al., 2018; Spackman et al., 2015), and a low dose was sufficient for H9N2 UDL/08 (Clements et al., 2020) to cause productive infections, we inoculated chickens with a high virus dose mix of Anhui/13 and UDL/08 which included a predominance of the former. The observed virus shedding from directly infected chickens indicated that a productive infection was established. All chickens placed in contact with the infectious birds also became infected and had a similar virus shedding profile to the directly inoculated chickens. As evident from the total AIV shedding in the directly- and contact-infected chickens, the reassortant viruses were shed at higher levels from oropharynx compared to cloaca. Contact transmission of infection indicated that fit viral progeny of unknown genotype (or genotype combinations) was produced following the mixed inoculation. Initial screening of total RNA extracts from oropharyngeal swabs by the genotyping RT-qPCRs inferred the presence of possible Anhui/13 and UDL/08 reassortants.

The majority of reassortant viruses derived their haemagglutinin (HA) and polymerase (PB2, PB1 and PA) genes from H9N2 UDL/08 and the neuraminidase (NA) gene from the H7N9 Anhui/13 virus. An analysis of polymerase activity of H9N2 and H7N9 based on *in vitro* reporter gene expression showed that H9N2 RNP complex had more than 200% activity compared to H7N9 RNP complex in chicken cells. This higher polymerase activity suggests a functional advantage which would have favoured the generation of reassortant H9N9 viruses containing polymerase genes from H9N2 viruses in chickens (Li et al., 2008; Octaviani et al., 2011).

The process of reassortment may either lead to attenuation of influenza A viruses in their hosts or may selectively increase the viral fitness (Zaraket et al., 2015) due to ‘genetic tuning’ of different gene segments (Wang et al., 2014). The multi-step replication kinetics of reassortant viruses indicated that viruses which had the M gene from Anhui/13 H7N9 showed increased replication in avian cells as reported earlier (Hao et al., 2019). Five reassortant H9Nx viruses, namely genotypes 118, 119, 121, 122 and 124 were more lethal in embryonated chicken eggs compared to the progenitor H7N9 and H9N2 viruses. This increased mortality in chicken embryonated eggs could be partly due to increased replication in avian cells which was significant for genotypes 119, 121 and 126, and was also observed for genotype 122 (although not significant). Four reassortant viruses which had the M gene derived from Anhui/13 (genotypes 119, 121, 123 and 126) showed increased replication in MDCK cells compared to UDL/08, while Anhui/13-origin M-gene in an H9N2 genetic backbone has been shown to increase virulence in a mouse infection model (Bi et al., 2015).

IAV co-infection *in vivo* can generate viral progenies with incomplete viral genomes (IVGs) which cannot replicate autonomously (Brooke et al., 2014). As such, plaque purification from unextracted swabs followed by the genotyping RT-qPCRs was done to identify viable reassortants. Lesser genotypes of reassortant viruses were identified after plaque purification compared to when swab samples containing a reassortant mix were directly subjected to genotyping by RT-qPCR. These findings suggested that total viral particle: infectious viral particle (PFU) ratio was high *in vivo* with only a minority of viruses capable of establishing productive infection at limiting dilutions in plaque assays as observed previously (Brooke, 2014; Donald and Isaacs, 1954; Enami et al., 1991; McLain et al., 1988; Noton et al., 2009; Wei et al., 2007). Further, the naked viral RNA picked up while swabbing could also have been detected by RT-qPCR. The exact constellation of segments in viruses isolated from the oropharyngeal swabs of contact chickens identified *bona fide* reassortants, capable of independent replication and *in vivo* transmission in chickens. To select a reassorted genotype for further evaluation with respect to zoonotic potential, the assessment was done by identifying the % genotype frequencies in the swab samples and all the highly abundant genotypes were then compared for their replication kinetics in human A549 cells. The H9N9 (genotype 122) was chosen as a viable reassortant which successfully emerged *in vivo* from infected chickens as a relatively abundant genotype having greater embryo lethality in chicken eggs and having more dynamic replication in A549 cells, and therefore selected as the likely candidate for assessing zoonotic potential.

For an AIV to cross the species barrier and infect humans requires an adaptive change which includes a shift in binding preference of viral glycoproteins towards human receptors (6SLN) (Herfst et al., 2012) and an increased stability reflected in a lower pH of endosomal membrane fusion (Imai et al., 2012). The receptor binding preference of reassortant H9N9 virus (genotype 122) showed a significantly stronger binding for human 6SLN and avian 3SLN sialoglycans compared to the parental H9N2 virus, but its strongest binding avidity was towards the avian-like 3SLN(6-su) receptor analogue. Compared to the H7N9 parental virus, the binding preference of the genotype 122 H9N9 virus was comparable for the 6SLN but weaker for 3SLN sialoglycans, but the virus again showed its strongest binding towards the 3SLN(6-su) receptor analogue. Earlier analysis of N9 NA showed that haemadsorption sites could be responsible for increasing the overall avidity of the virus towards the sialoglycans (Benton et al., 2017; Uhlendorff et al., 2009). Thus, increased viral replication of reassortant viruses bearing the NA (N9) of Anhui/13 origin in human A549 cells and MDCK cells may be a consequence of the N9 enhancing the binding avidity to the human-like 6SLN receptors. Viruses with preferable binding towards 3SLN(6-su) may have an increased propensity for circulation in terrestrial poultry (Gambaryan et al., 2008; Peacock et al., 2020), and this observation was reflected in the successful generation and transmission of these H9N9 genotypes in chickens in our *in vivo* experiment. The H9N2 and genotype 122 H9N9 viruses were found to have an optimal pH fusion of 5.4, which was slightly lower compared to that of Anhui/13 H7N9 having an optimal pH fusion 5.6 as seen previously (Chang et al., 2020). The results suggested that the reassortant H9N9 virus has a relatively more acid stable HA and stronger avidity for both human-like (6SLN) and avian-like (3SLN and 3SLN(6-su) receptors, thereby maintaining adaptation of such viruses for poultry while also enabling additional zoonotic potential (Peacock et al., 2020; Russier et al., 2016).

Ferrets are widely recognised as an effective animal model for evaluation of virus transmission and pathogenesis in humans (Enkirch and von Messling, 2015). Thus, to assess the possible *in vivo* consequences of variable *ex vivo* receptor binding of a reassortant H9N9 virus, genotype 122 was assessed for infection and transmission in ferrets. This reassortant H9N9 virus was compared with one parental virus - H7N9 Anhui/13 virus for virus shedding, dissemination in internal organs and pathogenesis. Interestingly, the reassortant H9N9 virus showed similar peak titres to those of the H7N9 virus in the directly infected (D0) ferrets, although D0 ferrets infected with the H7N9 virus continued to shed viral components for a longer duration. The ferrets placed in direct contact (i.e. the R1_DC_ groups) with the H9N9 or H7N9 infected D0 ferrets also experienced viral shedding as evidence of successful transmission, although H9N9 shedding appeared to attain lower maximal levels and was of shorter duration compared to that in the H7N9 R1_DC_ group. Neither the H7N9 nor the H9N9 viruses transmitted to the ferrets in the indirect contact (R1_In_) groups, via the bio- aerosol route. The D0 and R1_DC_ ferrets in both the H7N9 and H9N9 infected groups seroconverted against the homologous viruses. Interestingly, among the R1_In_ ferrets, those in the H9N9 contact group also seroconverted, while the ferrets in H7N9 contact group did not. Although H9N9 virus was not detected in the R1_In_ ferrets by RT-qPCR, the seropositive findings suggested that H9N9 may have initiated a very limited or highly localised infection following respiratory droplet exposure. Such a restricted infection may have been below the sensitivity of RT-qPCR, but nevertheless elicited seroconversion. Ferrets infected with either H7N9 or reassortant H9N9 demonstrated that both of these viruses replicated (semi-quantitative gross, H&E and IHC pathologies) in the nasal cavity to a similar degree and caused lesions in the nasal mucosa (Supplementary Figure S4) and lungs (Supplementary Figure S5).

Sequence analysis of the variant H9N9 viruses from the directly in-contact ferrets, performed by next generation sequencing, revealed one non-synonymous amino acid polymorphism at the consensus level in the HA gene. This polymorphism resulted in an amino acid change from an Alanine to Threonine at amino acid position 180 (A180T in mature H9 peptide numbering; A190T in mature H3 peptide numbering). This amino acid change is in the 190- Helix proximal to the receptor binding site and has been shown to increase the binding avidity of H9N2 more than 3500-fold and 20-fold towards avian (3SLN) and human (6SLN) and receptors, respectively (Sealy et al., 2018). It has therefore been suggested that this mutation may be associated with mammalian adaptation (Sealy et al., 2018), and may have arisen to compensate for the stalk deletion present in the N9 protein (Gao et al., 2013) of the H9N9 reassortant in order to maintain the crucial HA-NA balance for successful viral entry and exit from cells during the infection cycle (Mitnaul et al., 2000).

The number of human infections associated with H7N9 virus in China have been reportedly reduced after implementation of extensive poultry vaccination during autumn 2017 (Zeng et al., 2018). However, the virus has reassorted with enzootic H9N2 viruses in Eastern China since 2014 (Han et al., 2014; Wang et al., 2014), leading to the emergence of H9N9 viruses in natural ecosystems bearing internal genes from Anhui/13 H7N9 during 2016-2019 (Bi et al., 2020). The reassortant H9N9 viruses with polymerase genes from Anhui/13 H7N9 were also identified by RT-qPCR from the cloacal swab samples taken from contact chickens in our study, but the cloacal samples were not processed for plaque purification of reassortant genotypes. Whether internal genes from Anhui/13 H7N9 can provide a fitness advantage, compared to the internal genes from G1 lineage H9N2 viruses, needs further investigation.

Collectively, our data show that co-circulation of H7N9 and H9N2 viruses of G1 lineage circulating in the Indian sub-continent and the Middle East could lead to the emergence of novel reassortant H9N9 viruses which can transmit in poultry with additional zoonotic potential. Further evolutionary adaptation could enable efficient transmission to and between mammalian species such as humans. Thus, co-circulation of H7N9 and H9N2 viruses in the same enzootic regions represent a credible pandemic threat, and therefore necessitates continuing vigilance, including the emergence of novel reassorted genotypes.

## METHODS

### Safety and Ethics Statement

The United Kingdom regulations categorise the H7N9 LPAIV as a SAPO 4 and ACDP hazard group 3 pathogen because it is a notifiable animal disease agent and presents a zoonotic risk (https://www.hse.gov.uk/biosafety/diseases/acdpflu.pdf). In addition, the necessary UK Genetic Modification guidelines were considered (https://www.hse.gov.uk/biosafety/gmo/acgm/acgmcomp/index.htm), hence all the *in vitro*, *in ovo* and *in vivo* experiments involving H7N9 virus and derivatives were done in licenced Containment Level 3 facilities at The Pirbright Institute (TPI) or Animal and Plant Health Agency (APHA). All animal studies and procedures were carried out in accord with the relevant United Kingdom and European regulations, and approval by the Animal Welfare Ethical Review Board (AWERB) at the APHA, Weybridge.

### Viruses used in the study

The nucleotide sequences of the different gene segments of H7N9 (A/Anhui/1/13, abbreviated to “Anhui/13”) and H9N2 (A/chicken/Pakistan/UDL-01/08, abbreviated to “UDL/08”) viruses were retrieved from Global Initiative on Sharing All Influenza Data (GISAID) webserver (https://www.gisaid.org/) and the different viral gene segments were synthesised by Geneart^TM^ (Thermo-Fisher Scientific) and subcloned into pHW2000 vector by standard cloning technique involving *BsmBI* sites (Hoffmann et al., 2001) or restriction enzyme and ligation independent technique (Bhat et al., 2020). The isolate IDs of the IAVs used in the study are listed (Supplementary Table S3).

### Cell culture

The Madin Darby Canine Kidney (MDCK), human embryonic kidney (HEK) 293T, chicken DF-1 and adenocarcinomic human alveolar basal epithelial cells (A549) (obtained from Central Services Unit at TPI) were maintained in Dulbecco’s Modified Eagle’s medium (DMEM) (Sigma), supplemented with 10% foetal bovine serum (FBS) (Life Science Production) and 1x penicillin streptomycin (Gibco). Primary chicken kidney (CK) cells were prepared as previously described (Penzes et al., 1994), and were maintained in EMEM (Sigma) containing 7% BSA (Sigma), 1x penicillin streptomycin (Gibco) and tryptose phosphate broth (Sigma). All cell lines and primary cells were maintained at 37℃ and 5% CO_2_.

### Virus rescue and propagation

The viruses were rescued by reverse genetics (RG) (Hoffmann et al., 2002; Hoffmann et al., 2000), confirmed by the haemagglutination (HA) assay using standard methods and propagated in 10 day old specific pathogen free (SPF) embryonated chicken eggs at 37°C for 72 hours (OIE, 2018a). The viruses were aliquoted and stored at -80°C until required when they were diluted appropriately in DMEM medium (Invitrogen) for *in vitro* infections or sterile phosphate buffered saline (PBS) for *in ovo* and *in vivo* infections.

### Quantification of viral inocula for *in vivo* experiments

Ten-fold serial dilutions of the H7N9 and H9N2 RG viruses were made and 100µl of each dilution was inoculated in a group of 6 embryonated chicken eggs. These were incubated at 37°C for 72 hours. Allantoic fluid was harvested from all the inoculated eggs and tested for the presence or absence of virus by the HA test (OIE, 2018a). Egg infectious dose 50 (EID_50_) titres were calculated by the method described by Reed and Muench (Reed and Muench, 1938).

### Experimental design: Co-infection of chickens, transmission

Twenty-seven SPF derived Rhode Island Red chickens (procured from the National Avian Research Facility (NARF), Roslin Institute, UK) were wing bled and swabbed (oropharyngeal and cloacal) prior to the commencement of the *in vivo* experiments. In order to exclude prior or ongoing IAV infection, the serum samples were tested by the Influenza A Antibody ELISA (IDEXX) which detects antibodies to the type-common NP antigen, and the swab samples were tested by M-gene RT-qPCR (detailed below). The transmission study included nine of these chickens at three weeks of age, referred to herewith as the D0 (“donor”) chickens, which were directly infected by intranasal (i.n.) inoculation with 200µl of mixed inoculum containing 1X10^5^ EID_50_ of H9N2 UDL/08 and 1X10^8^ EID_50_ of H7N9 Anhui/13. An equal number of age-matched chickens, referred to herewith as the R1 (“recipient”) chickens, were introduced at 1-day post-infection (dpi) for co-housing to serve as transmission contacts. Two other groups (n=3 chickens per group) served as singly infected control groups for i.n. inoculation with 200 µl of the same doses of the individual H9N2 or H7N9 RG viruses. In order to investigate the pathogenesis (organ tropism) of the mixed viral infection, another group of six age-matched chickens were housed separately and similarly infected via the i.n. route with the H7N9/H9N2 mix and were dedicated for pre-planned culling of three chickens at 2 and 4 dpi for post mortem (PM) analysis. Oropharyngeal and cloacal swabs were collected daily from all the D0 and R1 chickens in the transmission study and the singly infected controls until 14 dpi, and from the six chickens which were pre-planned for culling in the pathogenesis experiment. Where relevant, reference is made to the R1 chickens at “days post contact” (dpc) which corresponded to one day less than the dpi. All swabs were processed in 1ml WHO virus transport media (VTM) (WHO, 2006) and stored at -80℃ until further use. The chickens were monitored twice daily for clinical signs, and all were humanely euthanized by an overdose of pentobarbitone and terminally heart bled at the end of the transmission and pathogenesis experiments.

### Avian influenza virus RT-qPCRs

Viral RNA was extracted from the swab samples and tissue homogenates by robotic and manual methods, respectively (James et al., 2019a). For initial screening purposes, all extracted RNA samples were tested by RT-qPCR using primers and probes specific for M gene (Nagy et al., 2010) as described previously (Slomka et al., 2018). A ten-fold dilution series of RNA extracted from the titrated H7N9 and /or H9N2 (known EID_50_ titre) virus was used to plot a standard curve along with the positive threshold at Ct 36. To characterise the subtype among the progeny viruses generated after co-infection, RT-qPCRs for H7, H9, N2 and N9 genes were performed as previously (James et al., 2019a; Slomka et al., 2013; Slomka et al., 2009). For the six internal gene segments, the genotype was further characterised by using gene-specific primer and probes which distinguished the origin of the genetic segment of interest, i.e. Anhui/13 or UDL/08 which were detected by FAM or HEX fluorescence, respectively. The primer and probe details are listed in Supplementary Table S1.

### Plaque assay and plaque purification of viruses

To identify viable reassortant viruses that emerged after co-infection of chickens, plaque purification of virus from the contact oropharyngeal swab samples was carried out in MDCK cells. The MDCK cells in six-well plates were infected in quadruplicate with a ten-fold serial dilution of swab sample in a 500µl volume and incubated for 1 hour at 37°C. The virus inoculum was removed and 2ml of overlay media containing 2% agarose was added at 37°C and allowed to set. The plates were inverted and incubated for 4 days at 37°C, or until visible plaques were formed. Discrete plaques were harvested individually into 200µl of plain DMEM medium using a pipette tip. RNA was extracted from the media plaque suspensions by robotic methods and genotyped by RT-qPCR. For the *in vitro* experiments, all the viruses were titrated as plaque forming units (pfu)/ml.

### Multi-step replication kinetics of viruses

The reassortant viruses which were shed from contact chickens after co-infection were identified by RT-qPCR and rescued *in vitro* by RG. The replication kinetics of the reassortant RG viruses was assessed in CK, MDCK and A549 cells, as previously described (James et al., 2019b). CK and MDCK cells were infected with 0.0002 multiplicity of infection (moi) and A549 cells were infected with 0.05 moi of respective viruses in infection medium (DMEM containing 1 x penicillin streptomycin and 0.3% BSA). The cell supernatant from four biological replicates was harvested 24, 48 and 72 hrs post-infection, and titrated by plaque assay.

### Minireplicon assay

Polymerase activity was assessed *in vitro* by plasmid based reporter gene expression as previously (te Velthuis et al., 2018). Chicken DF-1 cells and human HEK-293T seeded in 24- well plates were transfected with expression plasmids for different RNP combinations prepared from both the Anhui/13 and UDL/08 progenitor viruses by using Lipofectamine 2000 (Invitrogen), according to the manufacturer’s recommendations. 80 ng of PCAGGS plasmid encoding PB2 and PB1, 40 ng of PCAGGS plasmid encoding PA and 160 ng of PCAGGS plasmid encoding NP were co-transfected with 40 ng of a *Renilla* Luciferase pCAGGS expression plasmid and 80 ng of a pCk-PolI-Firefly plasmid expressing negative-sense firefly luciferase flanked by non-coding region of NS under the control of chicken specific polymerase I promoter (Long et al., 2016). The cells were incubated for 24 hrs at 37℃ (for HEK-293T) or 39℃ (DF-1) and lysed with 100µl of 1X passive lysis buffer. The luciferase activity in the transfected cells was measured by using a Dual Glo luciferase assay system (Promega). The polymerase activity was calculated by normalising firefly luciferase activity relative to the *Renilla* luciferase activity. The percent relative polymerase activity (% RPA) was calculated relative to the progenitor H7N9 or H9N2 positive controls, while negative controls excluded the PB1 plasmid during transfection.

### Virus purification and biolayer interferometry

Virus purification and biolayer interferometry was performed as described previously (Sealy et al., 2018). Briefly, the embryonated egg-propagated IAVs were ultra-centrifuged at 135200xg for 2hrs, purified on a continuous 30%-60% sucrose gradient and resuspended in PBS. The purified viruses were quantified by solid-phase indirect ELISA (Supplementary methods) and tested in an Octet RED bio-layer interferometer (Pall ForteBio, California, CA, USA) for receptor binding against sialoglycopolymers - α2,3-α1-4(6-HSO_3_)GlcNAc (3SLN(6-su)), as sialyllactosamine (3SLN) or Neu5Ac described previously (Lin et al., 2012). Virus association with the bound receptor analogues was measured at 20 °C for 30 in. Virus-binding amplitudes were normalized to fractional saturation of the sensor surface and plotted against sugar loading. The relative dissociation constant (K_D_), as a measure of binding to 6SLN, 3SLN and 3SLN (6Su), was calculated.

### pH stability of viruses

The acid-stability (Russell, 2014) of the selected reassortant H9N9 (genotype 122) and parental H7N9 and H9N2 viruses was determined by the ability of reassortant viruses to form syncytia in infected Vero cells exposed to different pH conditions. The viruses diluted two-fold in infection medium (DMEM containing 1x penicillin streptomycin) were used to infect Vero cells in 96 well format. The highest virus dilution infecting 100% of the Vero cells was calculated by immunostaining (Supplementary methods). This viral titre was used to infect Vero cells in 96-well-plates for the syncytium formation assays. At 16 h post infection, μg/ml TPCK trypsin for 15 min and then exposed to PBS buffers with pH values ranging from 5.2 -6.0 (at 0.1 pH-unit increments) for 5 min. The PBS buffer was then replaced with DMEM containing 10% FCS. The cells were further incubated for 3 h at 37 °C to allow for syncytium formation before being fixed with ice-cold (-20℃) methanol and acetone (1:1) mixture for 12 mins. and stained with 20% Giemsa stain (Sigma-Aldrich) for 1 h at room temperature. The pH at which syncytium formation was judged to be greater than 50% corresponded to the pH of viral membrane fusion.

### Experimental design: Ferret infection and transmission

Thirty male ferrets (*Mustela putorius furo)* were sourced from Highgate Farms, UK at a maximum age of 3 months and weighing between 750-1000g. The ferrets were confirmed as serologically negative to IAV by ID Screen® Influenza A Nucleoprotein Indirect ELISA (ID Vet). Ferrets were also confirmed negative for IAV ongoing infection (shedding) by testing RNA extracted from nasal washes by the M-gene RT-qPCR (Nagy et al., 2010), as above. All ferrets were microchipped (bio-thermal chip) to monitor identification number and the body temperature. Two groups of ferrets (n=3 per group; the D0 ferrets) were housed in cages in separate containment rooms, and directly-infected via the i.n. route with 1 X 10^7^ EID_50_ of Anhui/13 (H7N9) or H9N9 (genotype 122) (Supplementary Figure S7). At 1 dpi, direct-contact ferrets (n=3, i.e. the R1_DC_ ferrets) were introduced for co-housing in the same cage with the D0 ferrets in each room. Simultaneously, indirect-contact (R1_In_) ferrets (n=3) were housed in a cage adjacent to that which housed the D0 and R1_DC_ ferrets in each room (Supplementary Figure S7). Both cages in each room were separated by a double mesh which prevented direct contact between the ferrets but allowed potential IAV aerosol transmission. Each room also contained a third cage which housed three ferrets directly-infected with the two IAVs, and these were culled at 4 dpi for post mortem (PM) analysis, these six being referred to as the D0_PM_ ferrets. Tissues from the D0_pm_ ferrets were taken into 1ml of PBS (10% w/v) and RNA extracted for testing for influenza virus RNA using M-gene RT-qPCR. All directly-infected and contact-exposed ferrets were nasal washed with 1 ml of PBS (0.5 ml/nare) and similarly tested for IAV RNA using M-gene RT-qPCR until 12 dpi. The remaining ferrets were culled and cardiac bled at 12 dpi, with seroconversion to IAV assessed by the haemagglutination inhibition (HI) test using the homologous antigens, as described.

### Serology

To remove the non-specific inhibitors for HI, chicken sera were inactivated at 56 °C for 30 min, while ferret sera were incubated with 4 volumes of receptor destroying enzyme (APHA Scientific, Weybridge) for 1 hour at 37°C before being inactivated at 56 previously described (WHO, 2002). Seroconversion to the subtype-specific HA antigens was identified by HI assay (OIE, 2018a) using four haemagglutination units of homologous viruses as antigen. ID Screen® Influenza A Nucleoprotein Indirect ELISA, (ID Vet) and Influenza A Antibody ELISA (IDEXX) were performed according to manufacturers’ instructions.

### Histopathology and immunohistochemistry (IHC)

Formalin fixed tissues were processed by routine histology methods. Haematoxylin and eosin (H&E) staining and immunohistochemistry (IHC) against IAV nucleoprotein (NP) were performed on serially sectioned formalin-fixed paraffin-embedded tissues as previously described (Londt et al., 2008).

### Statistical analysis

Students t test and One-way ANOVA followed by Tukey’s multiple comparisons test was performed using GraphPad Prism version 8.00 for Windows, GraphPad Software, La Jolla California USA, www.graphpad.com. P value <0.05 was considered statistically significant.

## ACKNOWLEDGEMENTS

We would like to thank Dr Stephen Martin of The Francis Crick institute for use of his software for analysing the bio-layer interferometry results. We would also like to thank Alejandro Núñez, Carlo Bianco, Caroline Warren and Saumya Thomas for help processing samples.

## AUTHOR CONTRIBUTIONS

Conceptualization - SB and MI; Performed experiments – SB, JJ; PC, SW, JRS, JES, SM, APMB, BM, FL and HJE; Analyzed data – SB, JJ, MJS and MI; Manuscript, original draft – SB JJ; Edited the original draft – MI, MJS and SMB; Edited the final draft – all authors; Funding acquisition – MI

## SUPPLEMENTAL INFORMATION

### Supplementary methods

#### Immunostaining procedure - cells

Briefly, the viruses were serially diluted (two-fold) in DMEM containing 1x penicillin streptomycin to infect the Vero cells in a 96-well-plate for 1h. The inoculum was then removed and washed once with PBS before addition of DMEM medium with 5% FCS and 1x penicillin streptomycin for 16 h. The medium was removed, and cells were fixed with ice-cold methanol and acetone (1:1) mixture for 12 minutes at room temperature. The cells were washed once with PBS and blocked with blocking buffer (0.1% PBS tween containing 5% BSA) for 1hr at room temperature. The cells were incubated with anti-nucleoprotein (NP) mouse monoclonal antibody HB-65 [H16-L10-4R5 (ATCC® HB-65™)] [diluted 1:200 in dilution buffer (0.1% PBS-tween containing 0.5% BSA)] for 1 hour at room temperature. The cells were washed 4 times with wash buffer (0.1% PBS-tween) followed by incubation with horseradish peroxidase-labelled rabbit anti-mouse immunoglobulins (Dako, Denmark) (diluted 1:200 in dilution buffer). The cells were washed four times with wash buffer and developed using liquid DAB and substrate chromogen system (Dako, USA) as mentioned by the manufacturer.

#### Quantification of viruses using solid-phase indirect ELISA

The purified test viruses along with a reference virus [X-31 (reassortant virus carrying HA and NA from A/Aichi/2/68 and internal genes from H1N1/PR8] were diluted in carbonate bicarbonate buffer (pH 9.6) and coated on 96 well ELISA plates (Nunc MaxiSorp™) for overnight. The coated wells were permeabilized with 0.2% triton-X for 30 min at room temperature and then washed 4 times with wash buffer before incubating with blocking buffer for 1 hr. Each well was incubated with anti-NP mouse monoclonal antibody (HB65) (1:3000 dilution in dilution buffer) for 1hr. The plate was washed 4 times with wash buffer and then incubated with horseradish peroxidase labelled anti-mouse secondary antibody (Dako) (1:2000 dilution in dilution buffer) for 1 hr followed by addition of TMB substrate reagent set (BD OptEIA™). The reaction was stopped by addition of 1N H_2_SO_4_ and absorbance was measured at 450 nm. The concentration of the purified viruses was calculated by comparison of the estimated NP content of the reference virus X-31 as described elsewhere (Nicholson et al., 1998) (Lin et al., 2012) and expressed as picomolar (pM).

#### Next generation sequencing of viruses

Complete viral genome sequencing of the nasal wash sample of H9N9 infected ferrets was carried out MiSeq System (Illumina) as explained elsewhere (Puranik et al., 2020)

## Supplementary Figures

**Supplementary Figure S1.**
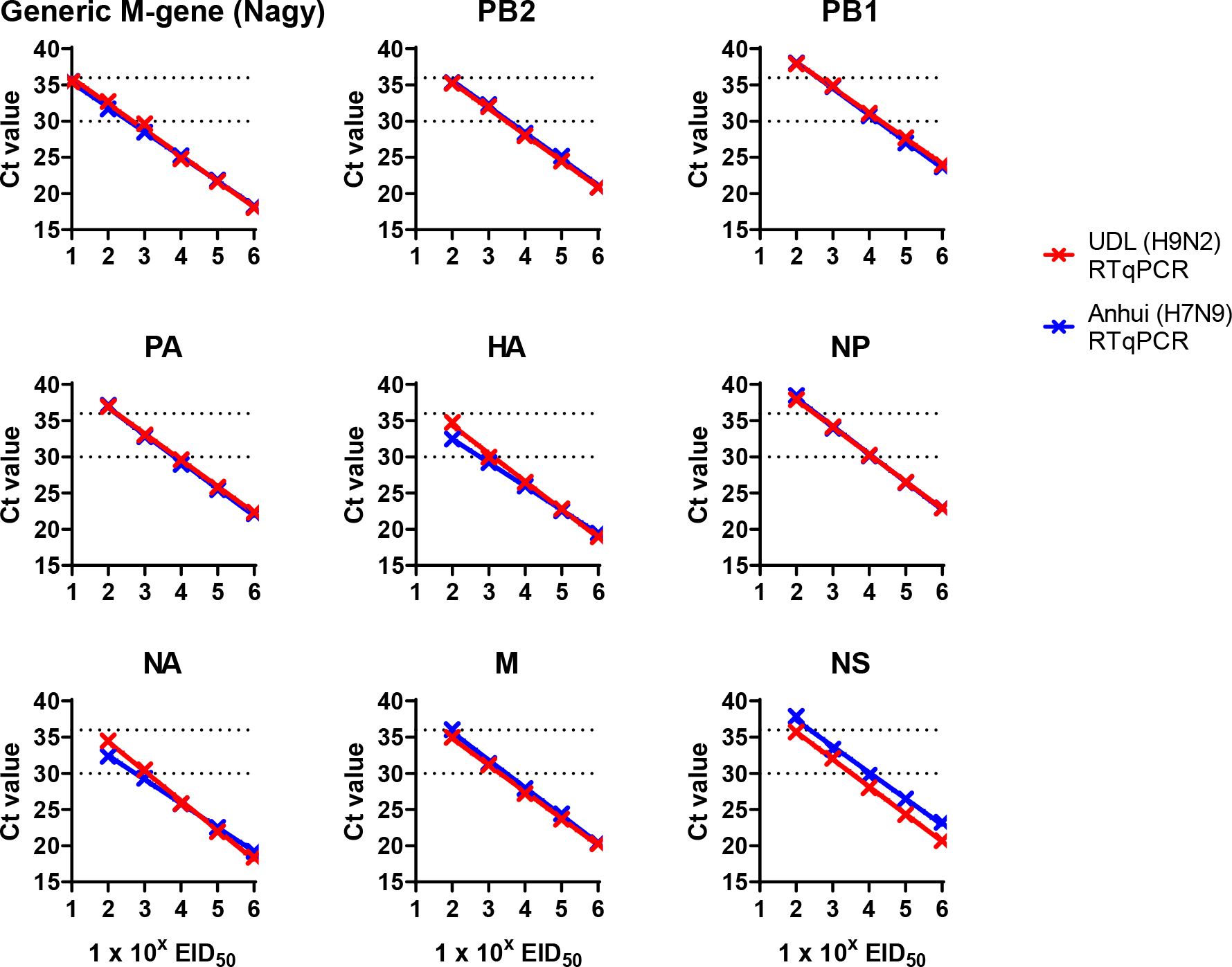
Segment specific RT-qPCR standard curves. RNA was extracted from both H7N9 Anhui/13 and H9N2 UDL/08 to achieve 1x106 EID_50_ / reaction. A 10-fold dilution standard curve was generated for each RNA and used in the gene specific RT-qPCR assays reactions. Ct values from each assay were plotted against EID50. The primer and probes designed to specifically detect H7N9 or H9N2 gene segment had comparable efficiency.

**Supplementary Figure S2:**
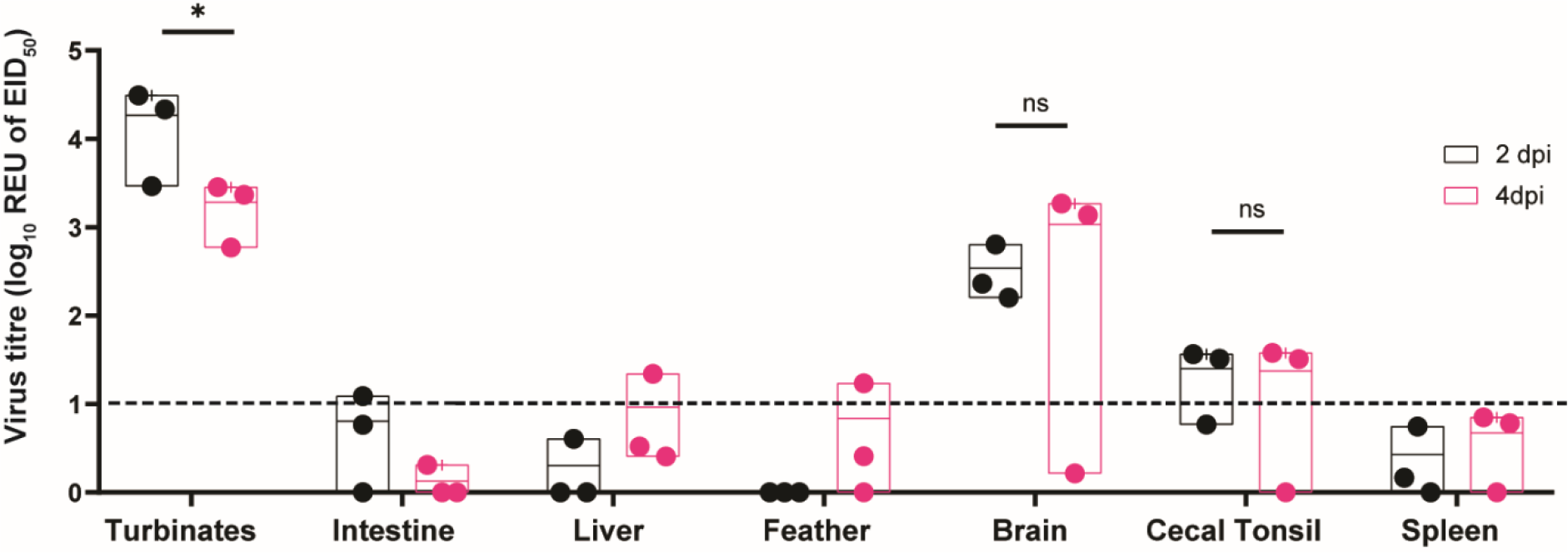
Virus dissemination in internal organs of chickens directly co-infected with Anhui1/13 H7N9 and UDL01/08 H9N2. Three co-infected chickens were pre-planned for cull at (A) 2dpi and (B) 4dpi. Virus dissemination in tissues was identified using AIV-generic M-gene primers and probes (Supplementary Table S1). The Ct values were compared against an Anhui1/13 or UDL/08 RNA standards to determine relative equivalency units (REU of EID_50_). The dotted line represents the positive cut-off REU value.

**Supplementary Figure S3:**
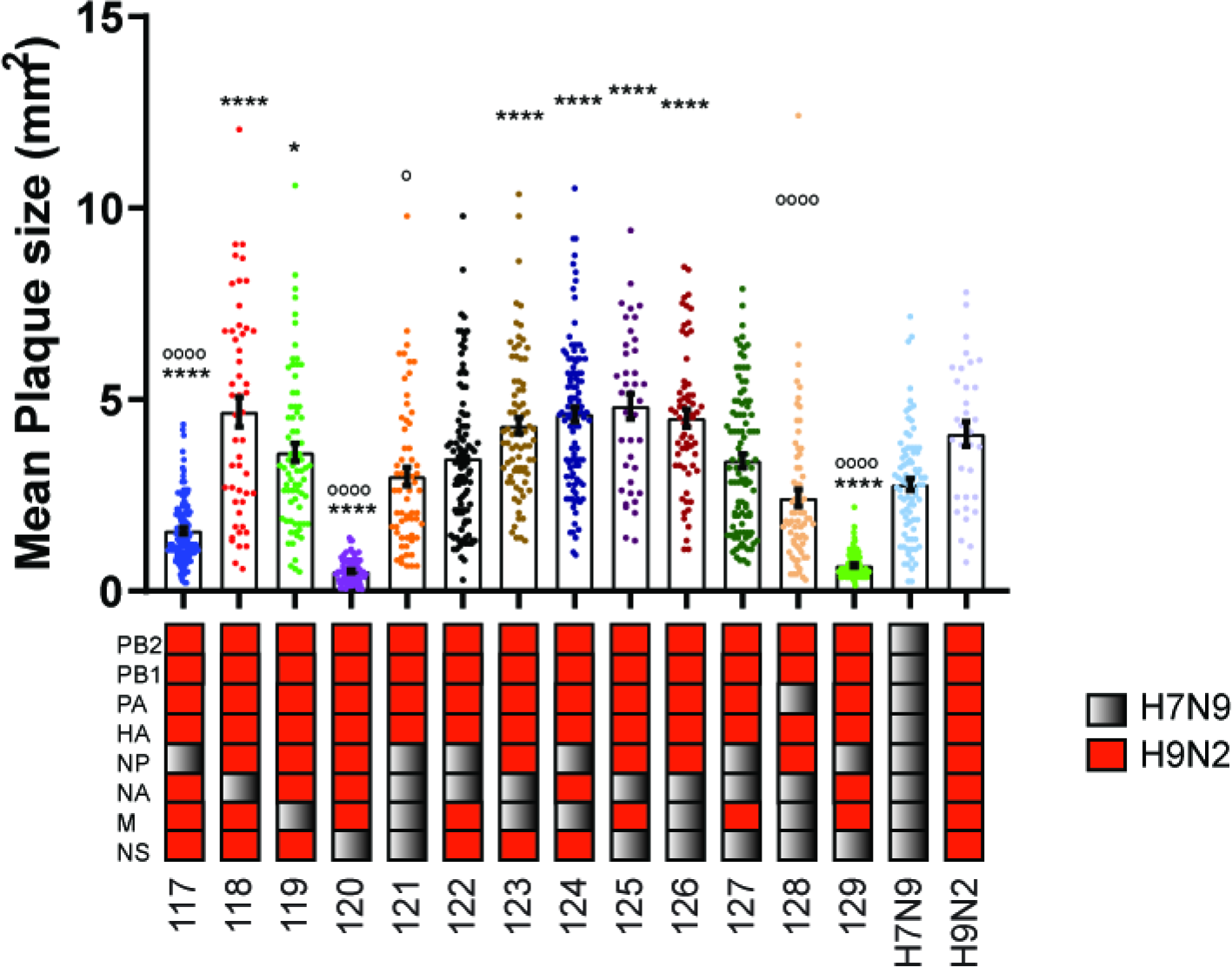
Mean Plaque size (mm2) of the reassortant H9Nx viruses on MDCK cells. Mean Plaque size (mm2) of the reassortant H9Nx viruses on MDCK cells. Plaque size was calculated in pixels using Image J software and converted to mm2. The mean plaque size of different reassortant H9Nx viruses was plotted along with Standard Error of Mean (SEM) and compared to H7N9 Anhui/13 virus and H9N2 UDL/08 virus. The mean plaque size of H9Nx viruses were compared to H7N9 virus using Dunnett’s multiple comparison test (one way ANOVA). The total number of counted plaques range from 33 - 125. As compared to the H7N9 virus, the plaque size of reassortant H9N9 virus genotype 117 (P<0.0001), genotype 118 (P<0.0001), genotype 119 (P < 0.05), genotype 120 (P<0.0001), genotype 123 (P<0.0001), genotype 124 (P<0.0001), genotype 125 (P<0.0001), genotype 126 (P<0.0001) and genotype 129 (P<0.0001) were significantly different. While as compared to H9N2 UDL/08 virus, the plaque size of genotype 117 (P<0.0001), genotype 120 (P<0.0001), genotype 121 (P<0.05), genotype 128 (P<0.0001) and genotype 129 (P<0.0001) were significantly different. ‘**** and *’ denote significance value of P<0.0001 and P < 0.05, respectively when compared to H7N9 virus. ‘oooo and o’ denote significance value of P<0.0001 and P < 0.05, respectively when compared to H9N2 virus.

**Supplementary Figure S4.**
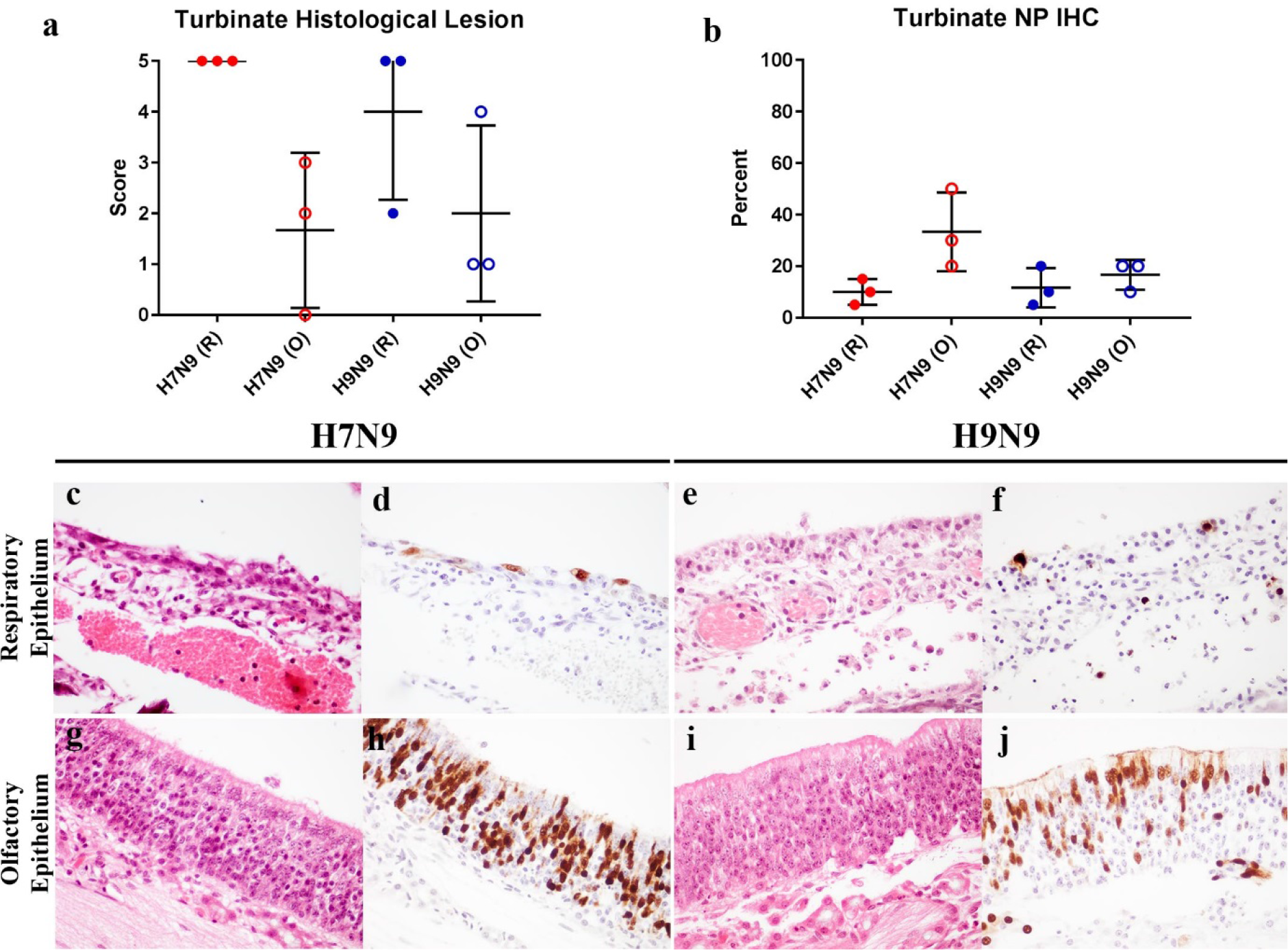
Ability of H7N9 or reasortant H9N9 IAV to infect and induce pathology in the nasal turbinates of D0_PM_ ferrets. Histopathological scores of nasal respiratory epithelium (R) and olfactory epithelium (O) **(a)**. Percentage of nucleoprotein (NP) immunopositive nasal epithelium (R or O) assessed by immunohistochemistry (IHC) **(b)**. Turbinate histological lesion and percentage of antigen labelling analysed with ANOVA with Tukey’s multiple comparison. Representative photomicrographs of ferret respiratory (**c-f**) and olfactory epithelial mucosa (**g-j**) assessed (left to right) alternately by H&E staining and NP IHC. Images originally produced at 400x magnification.

**Supplementary Figure S5.**
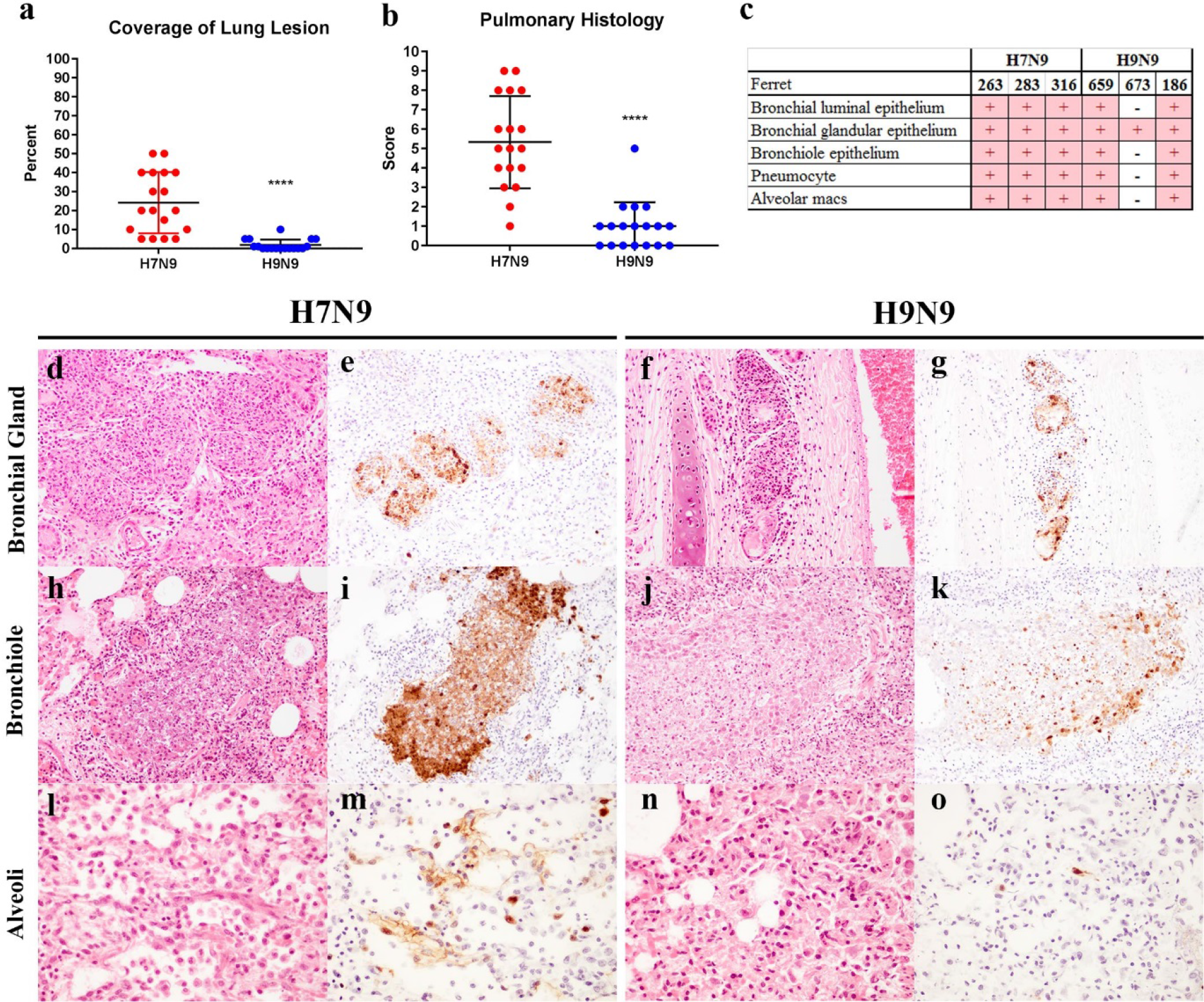
Ability of H7N9 or reasortant H9N9 IAV to infect and induce pathology in the lungs of D0_PM_ ferrets. Microscopy evaluation for area of lung lesions expressed in percentage **(a)** and pulmonary histopathology score **(b)**. Cellular distribution of IAV NP antigens in the lung as determined by immunohistochemistry (IHC), expressed as positive (+) or negative (-) **(c)**. Representative photomicrographs of ferret bronchial glands **(d-g)**, bronchiole **(h-k)**, and alveoli **(l-o)**. and olfactory epithelial mucosa **(g-j)** stained with H&E and NP IHC. Images taken at 200x magnification. Area of lung lesions and pulmonary histopathology score analysed with t-test. ****p<0.0001.

**Supplementary Figure S6.**
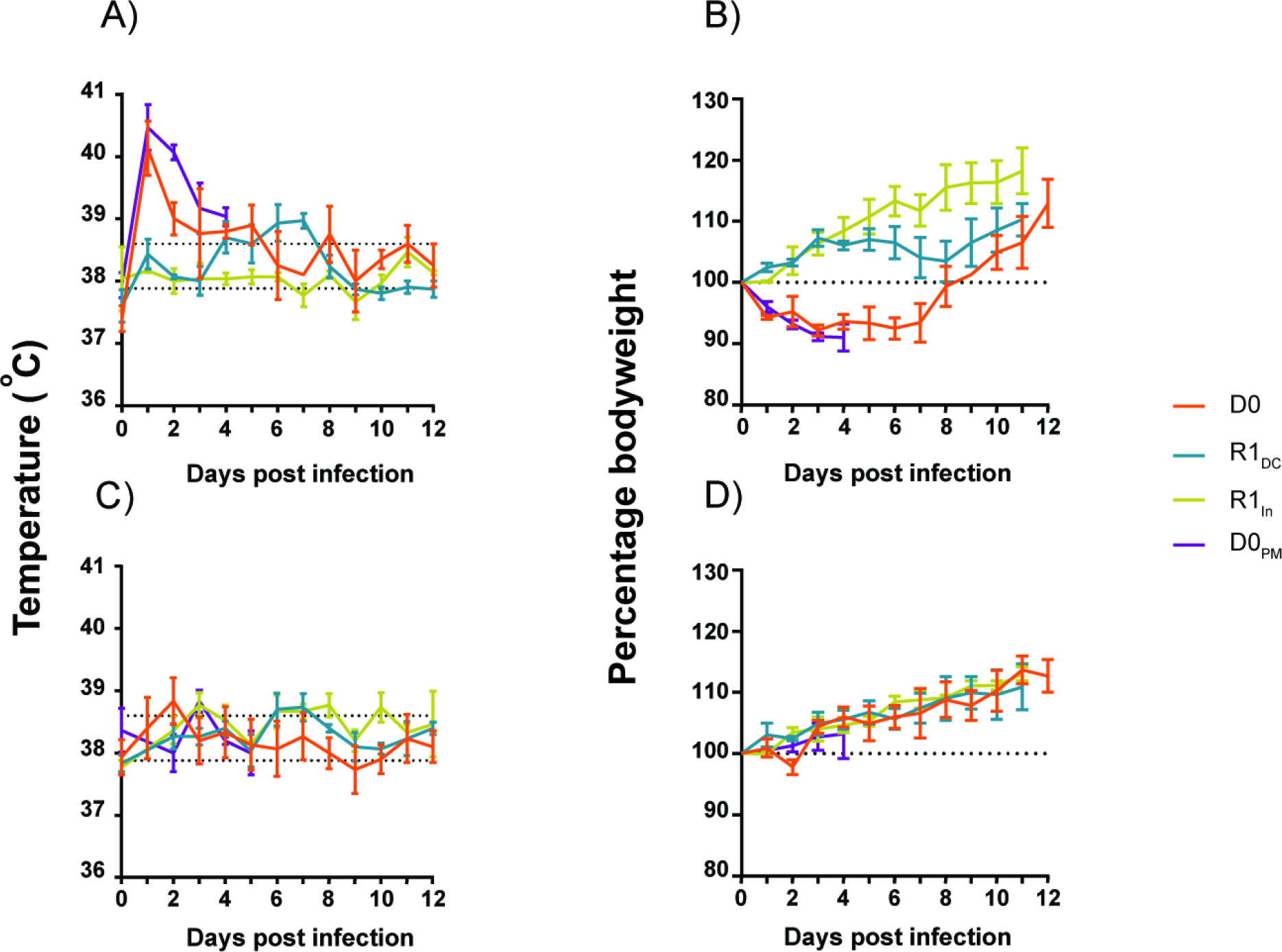
Changes in body temperature and body weight of ferrets infected with or exposed to Anhui/13 (A, B) and reassortant H9N9 (C, D) viruses. The infection or exposure was either through direct inoculation (D0), direct contact (R1_DC_) or indirect contact (R1_In_). **(A,C):** Ferrets were continuously monitored for body temperature after infection until 12 dpi / 11 dpc. The lower dotted horizontal baseline (37.8 °C) and upper dotted horizontal line (39.4 °C) indicates the normal ferret body temperature range. **(B,D):** The mean change in the body weight of the ferrets (shown as a percentage body weight) was monitored daily until 12 dpi / 11 dpc.

**Supplemetary Figure S7:**
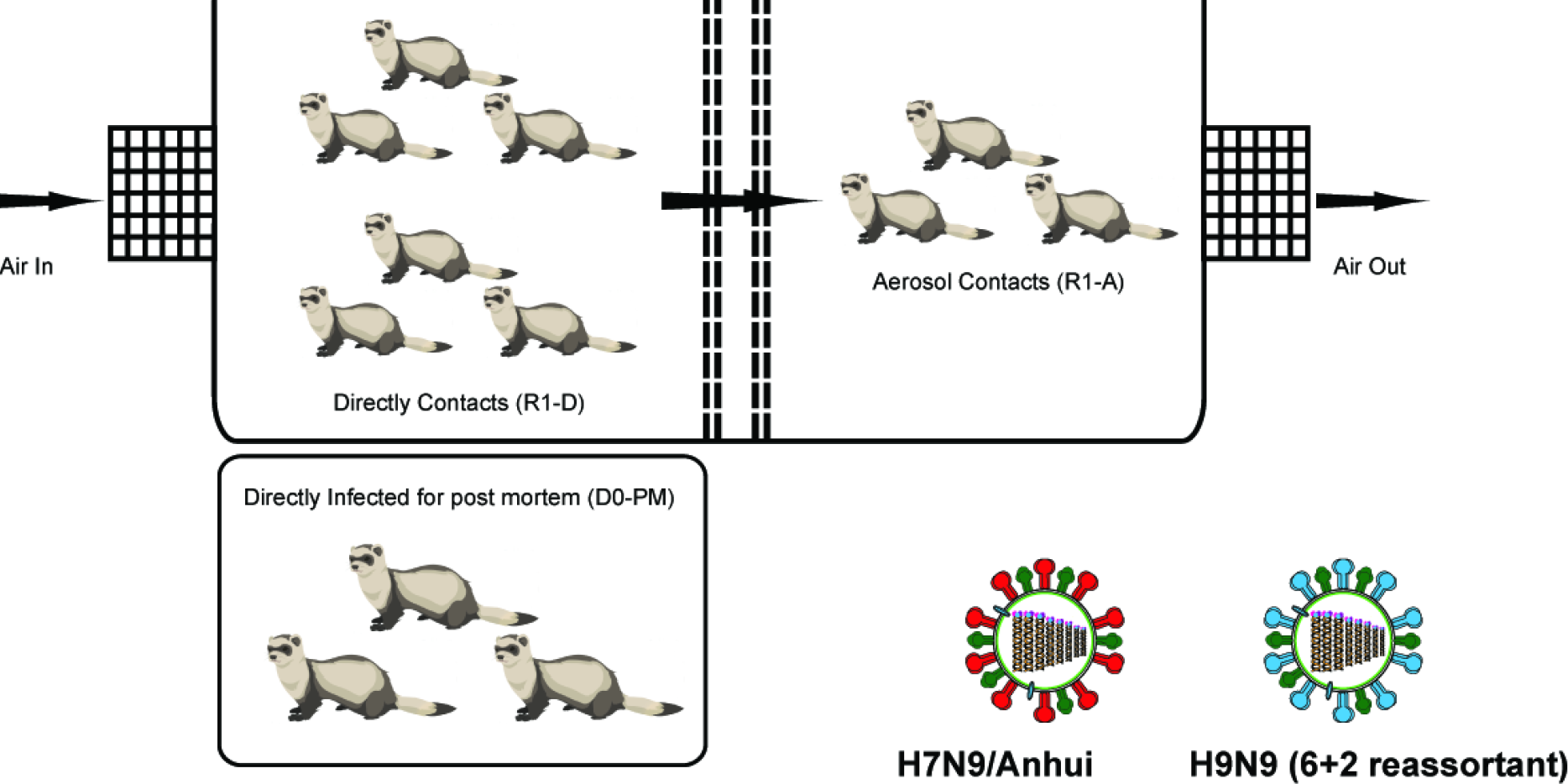
A schematic representation of ferret transmission experiment. Three ferrets were directly infected (D0) with H7N9 Anhui/13 virus or reassortant H9N9 virus. Contact ferrets (R1-D) were placed in contact with the directly infected ferrets in the same cage. A group of three ferrets were placed in an adjacent cage separated from the directly infected cage via a double mesh. The cages were maintained with a directional airflow from left to right (as shown by arrows above). A group of three directly infected ferrets (D0-PM) were culled on 4dpi to observe the virus dissemination in internal organs.

## Supplementary Tables

**Supplementary Table S1.**
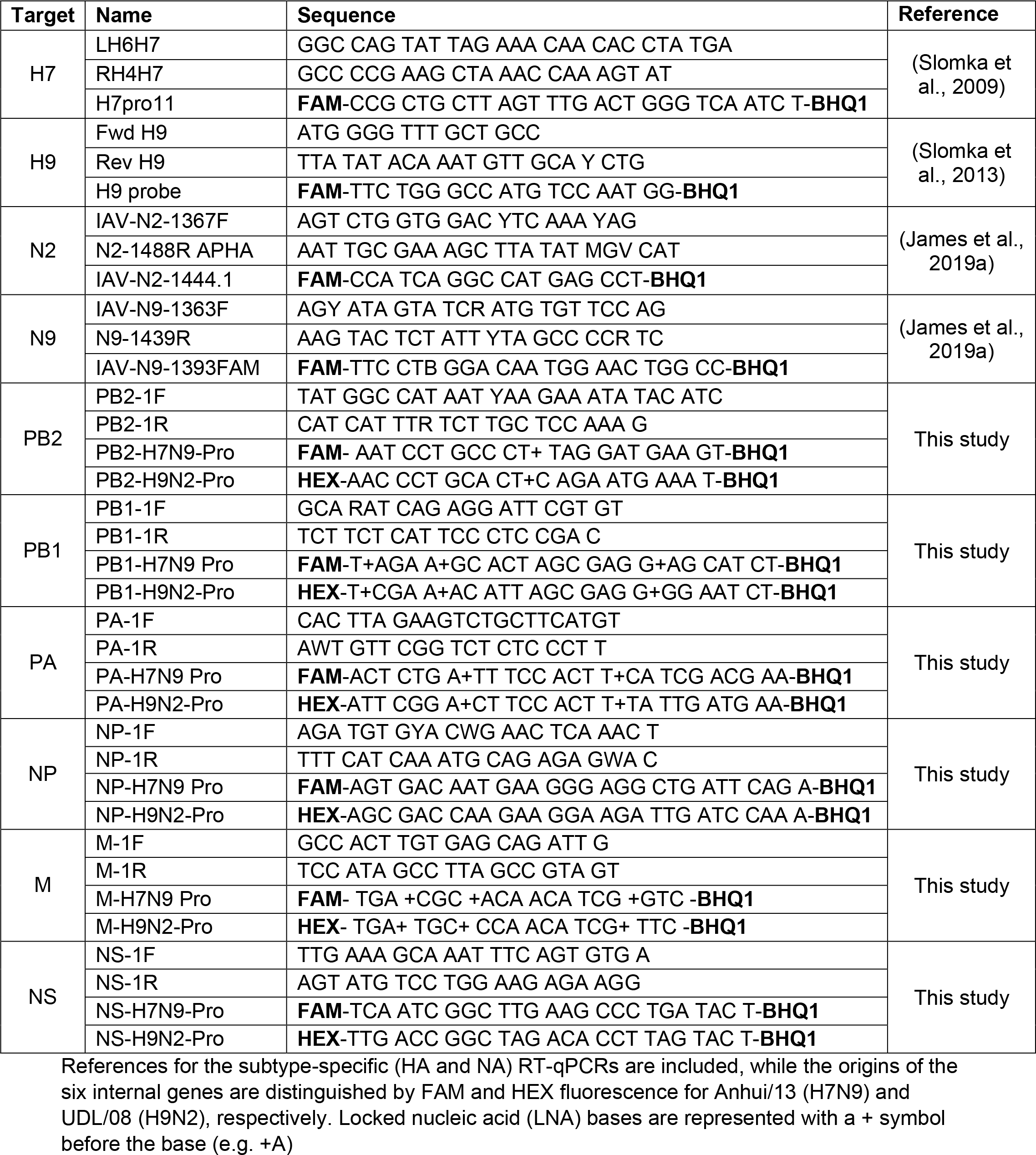
RT-qPCR primers and probes used in genotyping to identify the origins of the segments among the progeny viruses.

**Supplementary Table S2.** Distribution of virus antigens in th respiratory system of IAV infected ferret.

|  | H7N9 |  |  | H9N9 |  |  |
| --- | --- | --- | --- | --- | --- | --- |
|  | 263 | 283 | 316 | 659 | 673 | 186 |
| Nasal Turbinate | + | + | + | + | + | + |
| Cervical trachea | + | + | + | - | - | - |
| Thoracic trachea | + | + | + | - | - | - |
| Left cranial lung lobe | + | + | + | + | - | + |
| Left caudal lung lobe | + | + | + | + | + | + |
| Right cranial lung lobe | + | + | + | + | - | - |
| Right middle lung lobe | + | + | + | + | - | + |
| Right caudal lung lobe | + | + | + | + | - | + |
| Accessory lung lobe | + | + | + | + | - | - |

+ positive, - negative. Viral nucleoprotein antigen in the respiratory system of infected ferrets was detected by Immunohistochemistry (IHC) of different tissues.

**Supplementary Table S3.** Sequence accession numbers of the progenitor AIVs used in the present study

| S.No | Virus | Isolate ID |
| --- | --- | --- |
| 1 | A/chicken/Pakistan/UDL-01/2008 | EPI_ISL_29793 |
| 2 | A/Anhui/1/2013 | EPI_ISL_138739 |

The sequence of different gene segments of H9N2 UDL/08 and H7N9 Anhui/13 were retrieved from GISAID.

KEY RESOURCE TABLE.

KEY RESOURCE TABLE
| Reagent or Resource | Source | Identifier |
| --- | --- | --- |
| **Antibodies** |  |  |
| anti-nucleoprotein (NP) mouse monoclonal antibody | ATCC | HB-65[H16-L10-4R5 (ATCC® HB-65™)] |
| Polyclonal rabbit anti-mouse immunoglobulin HRPO | Dako | P0260 |
| **Bacterial or viral strains** |  |  |
| A/Chicken/Pakistan/UDL-01/2008/H9N2 | Iqbal Lab, TPI, UK | This study |
| A/Anhui/1/13/H7N9 | Iqbal Lab, TPI, UK | This study |
| Top 10 chemically competent E. coli | ThermoFisher Scientific | C404010 |
| **Commercial assays** |  |  |
| influenza A Antibody ELISA | IDEXX | 99-53101 |
| ID Screen® Influenza A Nucleoprotein Indirect ELISA | ID Vet | FLUNPS-10P |
| Dual Glo luciferase assay system | Promega | E2920 |
| **Experimental models: Cell lines** |  |  |
| Madin Darby Canine Kidney (MDCK) | Central services unit, TPI, UK | Not available |
| human embryonic kidney (HEK) 293T | Central services unit, TPI, UK | Not available |
| chicken DF-1 | Central services unit, TPI, UK | Not available |
| adenocarcinomic human alveolar basal epithelial cells (A549) | Central services unit, TPI, UK | Not available |
| Primary chicken kidney (CK) cells | Central services unit, TPI, UK | Not available |
| Vero cells | Central services unit, TPI, UK | Not available |
| **Experimental models: Animals/Strains** |  |  |
| Chickens/Rhode Island Red | Roslin Institute, UK | Not available |
| Specific pathogen free eggs | Valo Biomedica, GmbH | Not available |
| Ferrets | Highgate Farms, UK | Not available |
| **Oligonucleotides and probes** |  |  |
| Supplementary Table S2 | Iqbal Lab, TPI, UK | This study |
| **Recombinant DNA** |  |  |
| #875 PHW2000-PB2-UDL/08 | Iqbal Lab, TPI, UK | Clements et al., 2020 |
| #876 pollnewterm pMA-PB1-UDL/08 | Iqbal Lab, TPI, UK | Clements et al., 2020 |
| #914 pCAGGs- PB1 Pol2 Helper | Iqbal Lab, TPI, UK | Clements et al., 2020 |
| #874 PHW2000-PA-UDL/08 | Iqbal Lab, TPI, UK | Clements et al., 2020 |
| #872 PHW2000-HA-UDL/08 | Iqbal Lab, TPI, UK | Clements et al., 2020 |
| #871 PHW2000-NP-UDL/08 | Iqbal Lab, TPI, UK | Clements et al., 2020 |
| #873 PHW2000-NA-UDL/08 | Iqbal Lab, TPI, UK | Clements et al., 2020 |
| #869 PHW2000-M-UDL/08 | Iqbal Lab, TPI, UK | Clements et al., 2020 |
| #870 PHW2000-NS-UDL/08 | Iqbal Lab, TPI, UK | Clements et al., 2020 |
| #1091 pCAGGs-PB2-UDL/08 | Iqbal Lab, TPI, UK | Clements et al., 2020 |
| #600 pCAGGs-PB1-UDL/08 | Iqbal Lab, TPI, UK | Clements et al., 2020 |
| #1093 pCAGGs-PA-UDL/08 | Iqbal Lab, TPI, UK | Clements et al., 2020 |
| #1092 pCAGGs-NP-UDL/08 | Iqbal Lab, TPI, UK | Clements et al., 2020 |
| #924 PHW2000-NS-Anhui/13 | Iqbal Lab, TPI, UK | This study |
| #925 PHW2000-M-Anhui/13 | Iqbal Lab, TPI, UK | This study |
| #926 PHW2000-NP-Anhui/13 | Iqbal Lab, TPI, UK | This study |
| #927 PHW2000-PA-Anhui/13 | Iqbal Lab, TPI, UK | This study |
| #928 PHW2000-PB2-Anhui/13 | Iqbal Lab, TPI, UK | This study |
| #929 PHW2000-PB1-Anhui/13 | Iqbal Lab, TPI, UK | This study |
| #1002 pCAGGs-PB2-Anhui/13 | Iqbal Lab, TPI, UK | This study |
| #1003 pCAGGs-PB1-Anhui/13 | Iqbal Lab, TPI, UK | This study |
| #1004 pCAGGs-PA-Anhui/13 | Iqbal Lab, TPI, UK | This study |
| #1005 pCAGGs-NP-Anhui/13 | Iqbal Lab, TPI, UK | This study |
| pCAGGs Renilla Luciferase | Barclay Lab, Imperial college, London | Long et al., 2016 |
| pCk-Poll-Firefly | Barclay Lab, Imperial college, London | Long et al., 2016 |
| **Softwares** |  |  |
| Prism | GraphPad | Version 8.01 |
| **Cell culture media and Chemicals** |  |  |
| Dulbecco's Modified Eagle's medium (DMEM) | Sigma | D6429 |
| Foetal bovine serum (FBS) | Life Science Production | S-001A-BR |
| Minimum Essential Medium (MEM) -10X | Sigma | M0275 |
| Minimum Essential Medium (MEM) | Sigma | M2279 |
| Bovine Serum Albumin (BSA) | Sigma | A7906 |
| Penicillin-Streptomycin (10,000 U/mL) | Gibco | 15140-122 |
| Tryptose Phosphate Broth (TPB) | Sigma | T8159 |
| Virus Transport Medium (VTM) |  | WHO, 2006 |
| QIAmp viral RNA mini kit | Qiagen | 52904 |
| Low melting agarose | Oxoid | LP0028 |
| 2.5% trypsin | Gibco | 15090-046 |
| 1x dissociation medium (Versene) | Gibco | 15040-033 |
| Lipofectamine 2000 reagent | Invitrogen | 11668-019 |
| Opti-MEM™ I Reduced Serum Medium | Gibco | 31985-070 |
| Sodium bicarbonate | Sigma | S8761 |
| HEPES buffer | Sigma | H0887 |
| Sucrose | Sigma | S0389 |
| Neu5Aca3` (6-su)Lec-C3 | Lectinity | 0951-BP |
| 3`SLN-C3 | Lectinity | 0036-BP |
| 6`SLN-C3 | Lectinity | 0997-BP |
| Triton X-100 | Sigma | T8787 |
| Tween-20 | Sigma | P1379 |
| liquid DAB and substrate chromogen system | Dako | K346811-2 |
| TMB substrate reagent set | BD OptEIA™ | 555214 |
| Carbonate bicarbonate buffer | Sigma | C3041-50CAP |
| Receptor destroying enzyme (RDE) | APHA Scientific, UK | Not available |

